# SEDS-bPBP pairs direct Lateral and Septal Peptidoglycan Synthesis in *Staphylococcus aureus*

**DOI:** 10.1101/552034

**Authors:** Nathalie T. Reichmann, Andreia C. Tavares, Bruno M. Saraiva, Ambre Jousselin, Patricia Reed, Ana R. Pereira, João M. Monteiro, Rita G. Sobral, Michael S. VanNieuwenhze, Fábio Fernandes, Mariana G. Pinho

## Abstract

Peptidoglycan (PGN) is the major component of the bacterial cell wall, a structure essential for the physical integrity and shape of the cell. Bacteria maintain cell shape by directing PGN incorporation to distinct regions of the cell, namely through the localisation of the late stage PGN synthesis proteins. These include two key protein families, SEDS transglycosylases and the bPBP transpeptidases, proposed to function in cognate pairs. Rod-shaped bacteria have two SEDS-bPBP pairs, involved in cell elongation and cell division. Here, we elucidate why coccoid bacteria, such as *Staphylococcus aureus*, also possess two SEDS-bPBP pairs. We determined that *S. aureus* RodA-PBP3 and FtsW-PBP1 likely constitute cognate pairs of interacting proteins. Lack of RodA-PBP3 decreased cell eccentricity due to deficient pre-septal PGN synthesis, whereas the depletion of FtsW-PBP1 arrested normal septal PGN incorporation. Although PBP1 is an essential protein, a mutant lacking PBP1 transpeptidase activity is viable, showing that this protein has a second function. We propose that the FtsW-PBP1 pair has a role in stabilising the divisome at midcell. In the absence of these proteins, the divisome appears as multiple rings/arcs that drive lateral PGN incorporation, leading to cell elongation. We conclude that RodA-PBP3 and FtsW-PBP1 mediate lateral and septal PGN incorporation, respectively, and that the activity of these pairs must be balanced in order to maintain coccoid morphology.

---

Peptidoglycan (PGN) synthesis is an essential process that is both spatially and temporally regulated to ensure that the bacterial cell shape is maintained^1^. Rod-shaped bacteria elongate by synthesising PGN along the length of the cell in a process directed by the cytoskeletal protein MreB^2^. In *Escherichia coli* and *Bacillus subtilis*, this protein polymerises into short filaments that move processively around the cell diameter, and organise a multi-protein machinery, including PGN synthesis proteins, referred to as the elongasome or the Rod system^3-5^. Cell division is dependent on another cytoskeletal protein, FtsZ, which polymerises to form the Z-ring and recruits a multi-protein complex responsible for septum synthesis, known as the divisome^6,7^. This complex directs PGN incorporation to the midcell, resulting in inward PGN synthesis, and eventually bisects the mother cell, leading to daughter cell separation.

Ovococci such as *Streptococcus pneumoniae* and *Lactococcus lactis* lack MreB, and FtsZ is proposed to coordinate both elongation and septation^8,9^. In these organisms PGN is incorporated at the midcell in two defined modes: (i) at the lateral wall, eventually increasing the length of the cell and (ii) inwards, to synthesise the septum and future cell poles. Cocci, such as *Staphylococcus aureus*, also require FtsZ for divisome assembly and lack components of the elongasome, including MreB, consistent with its spherical shape^9^.

Directed PGN incorporation is accomplished by different sets of transglycosylases (TG) and transpeptidases (TP), the enzymes involved in the final stages of cell wall synthesis, that promote PGN polymerisation and cross-linking, respectively^10^. Key proteins involved in these steps are the penicillin-binding proteins (PBPs) that can be class A PBPs (aPBPs), with both TG and TP activity or class B PBPs (bPBPs) which possess only transpeptidase activity and have a second domain of unknown function^10^. Some bacteria also have monofunctional glycosyltransferases (MGTs), enzymes with a TG domain with homology to the TG domain of aPBPs^10^. More recently, RodA and FtsW, members of the shape, elongation, division and sporulation (SEDS) protein family, were also shown to have transglycosylase activity^11-13^. Together the SEDS and bPBPs have been proposed to form TG-TP cognate pairs, involved in cell elongation (RodA) and cell division (FtsW) in rod-shaped bacteria^11,14^. Compared to rod-shaped bacteria, *S. aureus* encodes a small number of PBPs: one bifunctional aPBP (PBP2), two bPBPs (PBP1 and PBP3) and one low molecular weight PBP (PBP4) with transpeptidase activity^15^. However, similarly to rods, it contains two SEDS proteins (FtsW and RodA). Given the proposed roles of SEDS-bPBP pairs in elongation and division of rod-shaped bacteria, why a coccus requires two sets of these proteins has remained a long-standing question.

Although the cell shape of *S. aureus* is close to spherical, we have recently used super-resolution microscopy to show that *S. aureus* cells elongate slightly during growth^16^. Here we investigated whether SEDS-bPBP cognate pairs occur in *S. aureus* and their role in directing PGN incorporation during the cell cycle.

## RESULTS

### RodA-PBP3 and FtsW-PBP1 form cognate pairs in *S. aureus*

To understand if SEDS proteins and bPBPs function together in cocci such as *S. aureus*, we investigated the essentiality of SEDS proteins FtsW and RodA, and bPBPs PBP1 and PBP3. Previous data has shown that PBP1 is essential^17^, while growth is unaffected in the absence of PBP3^18^. We found that we were able to delete *rodA* from *S. aureus* strain COL, but not *ftsW*. Therefore, *ftsW* was placed under the IPTG-inducible P*spac* promoter at an ectopic locus before deletion of the native copy, resulting in strain ColFtsWi. The same strategy was implemented for *pbpA* (encoding PBP1), giving rise to ColPBP1i. Growth curves of the null RodA (ColΔ*rodA*) and PBP3 (ColΔ*pbpC*) mutants mimicked that of COL, and the growth rate was maintained even in the absence of both genes (ColΔ*rodA*Δ*pbpC*) showing that RodA and PBP3 are not essential (Fig. 1a). In contrast, depletion of FtsW or PBP1 resulted in growth inhibition, indicating that these proteins are essential for cell viability (Fig. 1b). To determine if the essential functions of FtsW and PBP1 are related to their roles in PGN synthesis, the growth of active site mutants was assessed. Point mutations in conserved amino acids W121 and D287 of *S. aureus* FtsW, shown in *B. subtilis* RodA to abolish its transglycosylase activity^11^, were introduced into *S. aureus* FtsW-sGFP fusions expressed from a plasmid. Unlike wild-type FtsW-sGFP, these mutated proteins could not complement the growth of ColFtsWi in the absence of IPTG, although they localised to divisome, indicating that FtsW essentiality is dependent on its transglycosylase activity (Supplementary Fig. 1). In contrast, the PBP1 TP mutant carrying the S314A mutation in the transpeptidase active site, which had decreased PGN crosslinking, was not affected in terms of growth rate (Supplementary Fig. 2). This suggests that PBP1 has a second role, as previously proposed^19^. Compatible results have been obtained in *B. subtilis*, where introduction of a point mutation in the TP active site of the essential protein PBP2B, did not lead to loss of cell viability^20^.

**Figure 1.**
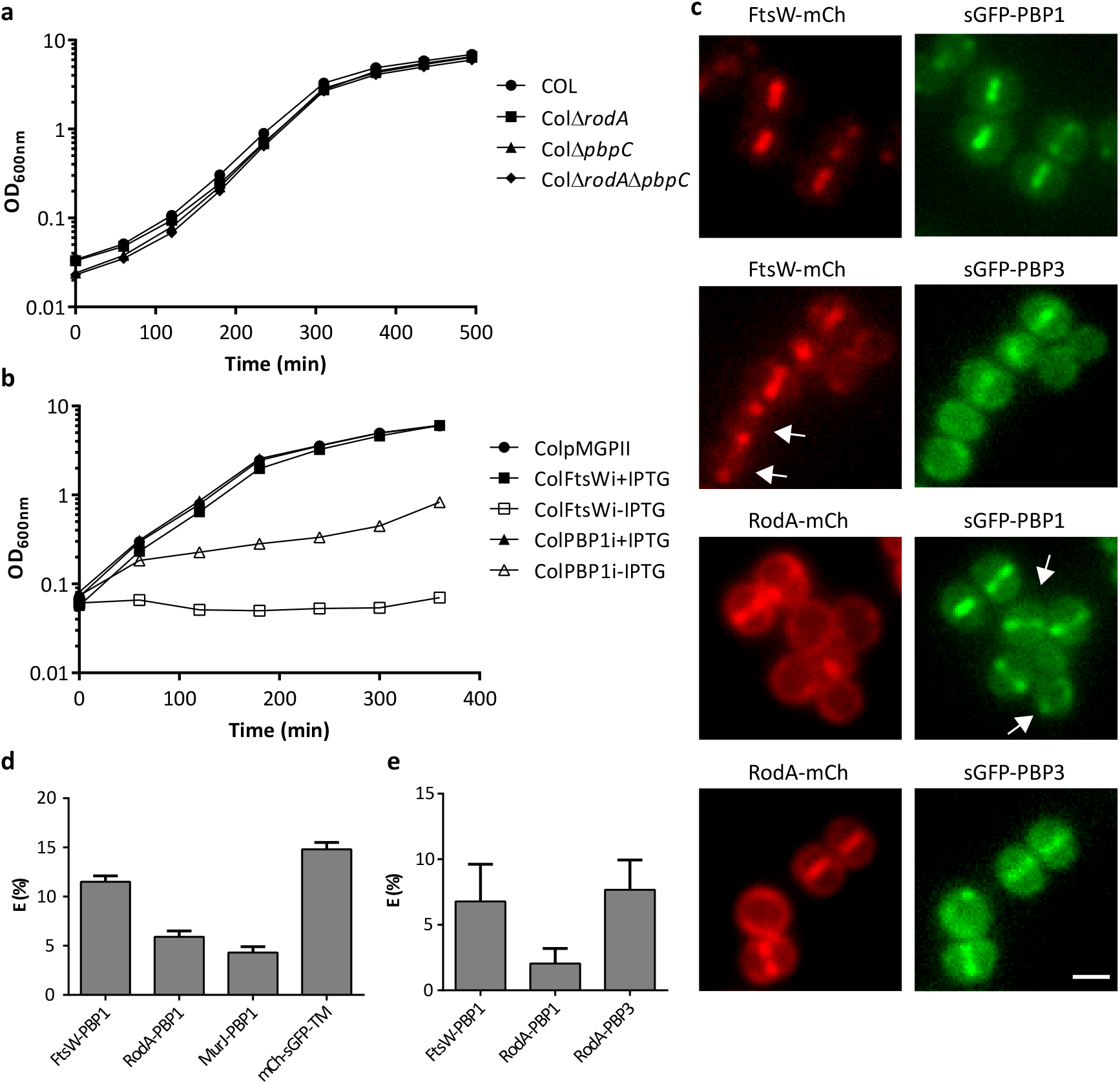
*S. aureus* has two cognate SEDS-bPBPs pairs of interacting proteins, FtsW-PBP1 and RodA-PBP3. a, Growth curve of parental strain COL and null mutants lacking RodA (ColΔ*rodA*) PBP3 (ColΔ*pbpC*) or both (ColΔ*rodA*Δ*pbpC*). **b**, Growth curve of strains expressing IPTG inducible FtsW (ColFtsWi) or PBP1 (ColPBP1i) grown in the presence or absence of IPTG. Control strain ColpMGPII was grown in the absence of IPTG. **c**, Epi-fluorescence images of strains ColWP1, ColWP3, ColP1pA and ColP3pA (from top to bottom) co-expressing different combinations of SEDS and bPBPs. Scale bar, 1 µm. Images are representative of three biological replicates. Arrows highlight cells with midcell localised FtsW but not PBP3, or midcell localised PBP1 but not RodA. **d**, FRET efficiency (E) determined by FLIM-FRET of strains ColpWP1, ColpAP1, ColpJP1 and ColpmCherry-sGFP-TM tandem control (where mCherry is linked to sGFP and anchored at the membrane). **e**, FRET efficiency (E) determined by seFRET of strains ColpWP1, ColpAP1 and ColpAP3, performed in triplicate. Data in panels d and e are represented as column graphs in which the height of the column is the mean and whiskers are s.d.

A recent study on the localisation patterns of PGN synthesis proteins demonstrated that FtsW and PBP1 are almost exclusively septal localised and arrive at the divisome at a similar time, while RodA and PBP3 are enriched at the septum but can also be observed at the peripheral membrane^21^. Given that cognate SEDS-bPBP pairs are likely to co-localise, we decided to correlate the time of arrival of each SEDS protein with the two *S. aureus* bPBPs in co-localisation studies. We observed cells with septal FtsW-mCh or sGFP-PBP1 but lacking septal sGFP-PBP3 or RodA-mCh, respectively (Fig. 1c, white arrows), whereas the reverse was not observed, demonstrating that FtsW and PBP1 arrive at midcell earlier than RodA and PBP3 (Supplementary Fig. 3).

Together, these results suggest that FtsW and PBP1 form an essential cognate pair that is recruited to the divisome before RodA and PBP3. To determine if these proteins directly interact, we performed Fluorescence Lifetime Imaging Microscopy - Fluorescence Resonance Energy Transfer (FLIM-FRET) using *S. aureus* cells expressing plasmid-encoded sGFP-PBP1 fluorescent fusion and mCherry fusions to the SEDS proteins. Since PBP1 localises to midcell, fluorescence lifetime analysis was restricted to septa. The co-expression of FtsW-mCh and sGFP-PBP1 in *S. aureus* cells caused a drop in the average fluorescence lifetime of sGFP in comparison to the lifetime of sGFP-PBP1 expressed alone, equating to a FRET efficiency (E) of 11.5 ± 0.6% (N=90) (Fig. 1d). Similarly high E values were observed for mCherry fused directly to sGFP and anchored to the membrane (tandem control mCh-sGFP-TM, E=14.8 ± 0.7%, N=50), while E values for RodA-mCh and sGFP-PBP1 were consistently lower (E=5.9 ± 0.6%, N=90) (Fig. 1d). These data indicate that PBP1 preferentially interacts with FtsW and not with RodA. To rule out the possibility of interactions being detected purely due to enrichment at the septum, an mCherry fusion to another almost exclusively septal localised protein, the putative lipid II flippase MurJ, was also probed for interactions with sGFP-PBP1, and was found to have low E values (4.3 ± 0.6%, N=90) (Fig. 1d). Despite attempts to detect interactions with the sGFP-PBP3 fusion by FLIM-FRET, its fluorescence was too weak under the conditions used. Therefore, we decided to test protein-protein interactions by sensitized emission FRET (seFRET), allowing lower expression levels of fluorescent fusions to be examined. Using this approach we were able to detect interactions between RodA-mCh and sGFP-PBP3 (E=7.7 ± 2.3%, N>45) (Fig. 1e). Furthermore, we detected a FRET efficiency of 6.8 ± 2.8% (N>45) for sGFP-PBP1 and FtsW-mCh, compared to 2.0 ± 1.2% (N>45) for sGFP-PBP1 and RodA-mCh (Fig. 1e), corroborating the FLIM-FRET data and providing *in vivo* evidence that both FtsW-PBP1 and RodA-PBP3 form cognate pairs.

### Elongation of *S. aureus* is mediated by RodA-PBP3

Having established that RodA-PBP3 and FtsW-PBP1 are the most likely *S. aureus* cognate TG-TP pairs, we decided to investigate their functions. *B. subtilis* RodA was recently shown to possess TG activity *in vitro* and its overexpression conferred resistance to moenomycin, an antibiotic that targets classical TG domains, but to which SEDS proteins are insensitive^11^. Given their essentiality, moenomycin sensitivity under FtsW or PBP1 depletion conditions could not be accurately determined. However, analysis of ColΔ*rodA* demonstrated that this strain was hyper-susceptible to moenomycin (Fig. 2a). Introduction of TG active site point mutations W111A and D286A (identified by Meeske and colleagues^11^) was sufficient to cause a decrease in minimum inhibitory concentration (MIC) without decreasing RodA-sGFP septal localisation (Fig. 2a,b). Although moenomycin does not inhibit TP activity, ColΔ*pbpC* had increased moenomycin susceptibility (Fig. 2a), indicating that RodA activity may be impaired in the absence of PBP3. Accordingly, we saw that RodA-sGFP requires PBP3 for correct septal localisation, while the opposite was not observed (Fig. 2b).

**Figure 2.**
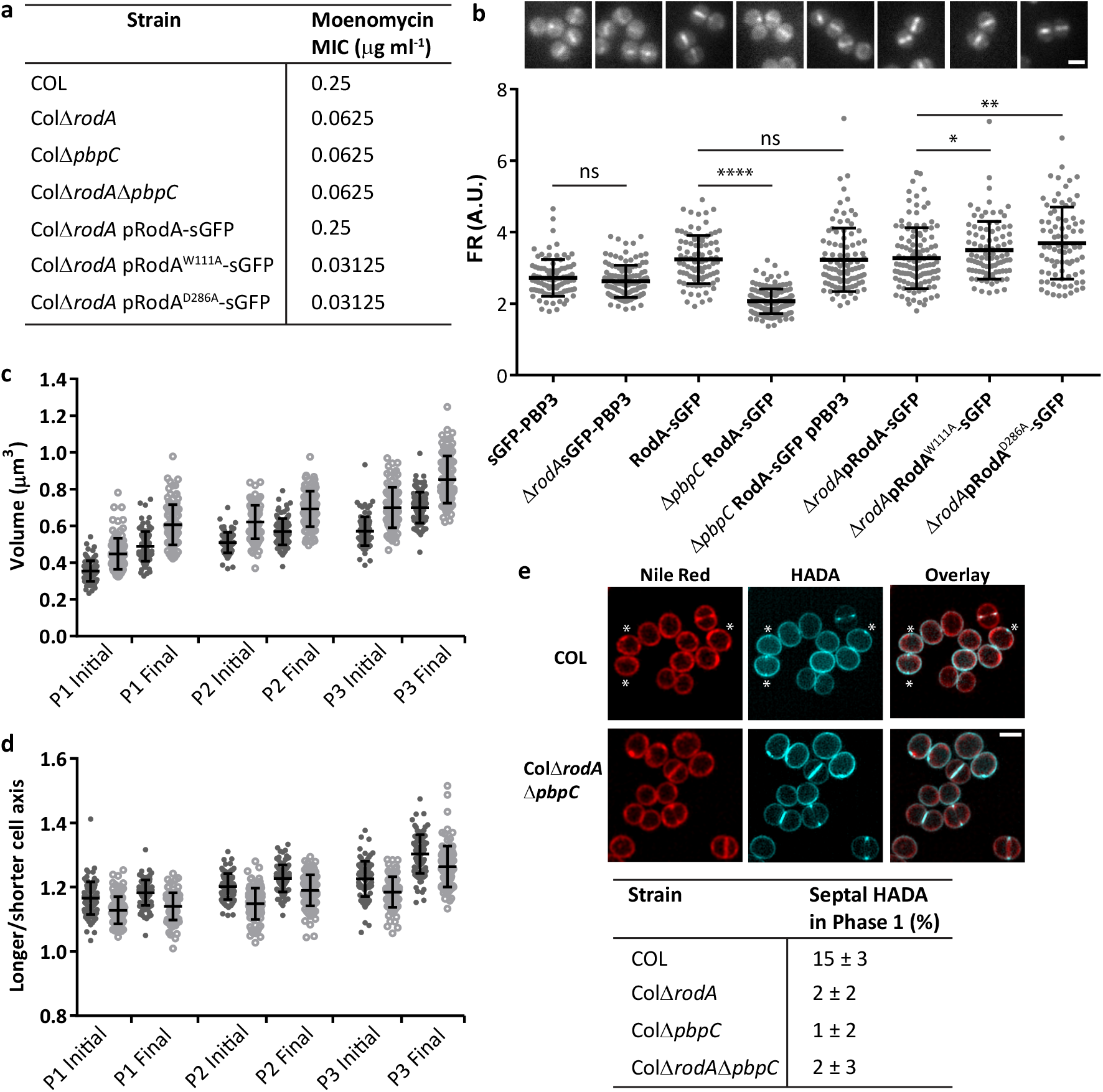
RodA-PBP3 have a role in cell elongation. a, Minimum Inhibitory Concentration (MIC) of moenomycin for (top to bottom) parental strain COL, null mutants lacking RodA, PBP3 or both and *rodA* null mutant complemented with sGFP derivatives of native or inactive RodA, performed in triplicate. **b**, Representative epi-fluorescence images (top) and Fluorescence ratios (FR, bottom) values for septal/peripheral signal in strains ColsGFP-PBP3, ColΔ*rodA*sGFP-PBP3, ColRodA-sGFP, ColΔ*pbpC*RodA-sGFP, ColΔ*pbpC*RodA-sGFPpPBP3, ColΔ*rodA*pRodA-sGFP, ColΔ*rodA*pRodA^W111A^-sGFP and ColΔ*rodA*pRodA^D286A^-sGFP. Scale bar, 1 µm. **c, d**, Cell volume (**c**) and cell axes ratios (**d**) of strain COL (filled dark grey circles) and ColΔ*rodA*Δ*pbpC* (open light grey circles) at the beginning and end of the three phases of *S. aureus* cell cycle. **e**, Structured Illumination Microscopy (SIM) images of COL and ColΔ*rodA*Δ*pbpC* stained with Nile Red (membrane) and HADA (PGN incorporation). Asterisks indicate cells with midcell HADA signal but no visible membrane invagination. Scale bar, 1 µm. Quantification of percentage of Phase 1 cells (cells with no septum) with midcell HADA signal in strains COL, ColΔ*rodA*, ColΔ*pbpC* and ColΔ*rodA*Δ*pbpC* (n=300). For panels b, c, d data are represented as scatterplots where the middle line represents the median and top and bottom lines show the standard deviation. Statistical analysis was performed using a two-tailed Mann-Whitney *U* test. * P<0.1; **P<0.01; \*\*\*\**P* < 0.0001. In panels c and d, *P* < 0.0001 for COL vs ColΔ*rodA*Δ*pbpC* was calculated in all cases.

Previous analysis of the cell shape of *S. aureus* has revealed that these cocci are slightly eccentric and elongate prior to septum synthesis^16^. We therefore evaluated whether a *S. aureus* SEDS-bPBP pair was responsible for this slight elongation. Although lack of RodA and/or PBP3 did not alter progression of the cell cycle (Supplementary Fig. 4), the double mutant was consistently enlarged, and cells were less elongated when compared to wild-type (Fig. 2c,d), pointing to a role of the RodA-PBP3 pair in generating the *S. aureus* slightly elliptical shape. For cell elongation to occur, PGN must be incorporated into defined regions of the cell. In rod-shaped bacteria, PGN synthesis is directed to the long axis by MreB, while ovococci (which lack MreB) incorporate PGN at the midcell along the lateral wall. To understand where PGN insertion takes place for *S. aureus* elongation, we used fluorescent D-amino acids (FDAAs)^22^ to label PGN synthesis. We found that 15 ± 3% (N=300) of the wild type cells had enriched HADA incorporation at midcell prior to any visible membrane invagination (Fig. 2e, asterisks), indicating that *S. aureus* elongation may occur through a process analogous to ovococci. Importantly, in the absence of PBP3 and RodA, the number of cells with this pre-septal HADA incorporation was reduced to less than 2 ± 3% (N=300), strengthening the notion that RodA-PBP3 mediates elongation of *S. aureus* by directing PGN synthesis to the lateral wall at midcell. Altogether, these data support a model whereby PBP3 recruits RodA to midcell and modulates its activity, and together this protein pair directs the lateral insertion of PGN that leads to elongation of the *S. aureus* cell.

### The FtsW-PBP1 cognate pair is essential for inward PGN incorporation at the septum

To determine the function of the essential proteins FtsW and PBP1, we imaged ColFtsWi and ColPBP1i depleted of IPTG over time. Both strains became progressively enlarged and with an indentation at midcell, clearly demonstrating a defect in cell division (Fig. 3a,b; Supplementary Figs. 5 and 6a,b). Most strikingly, and contrary to RodA and PBP3 deletion mutants, lack of FtsW or PBP1 led to elongated cells resembling rod-shaped bacteria (Fig. 3a,b; Supplementary Figs. 5 and 6a,b) not only highlighting the key role of FtsW-PBP1 in synthesising the septum, but also that the two cognate pairs most likely play opposing roles in defining the *S. aureus* cell shape.

**Figure 3.**
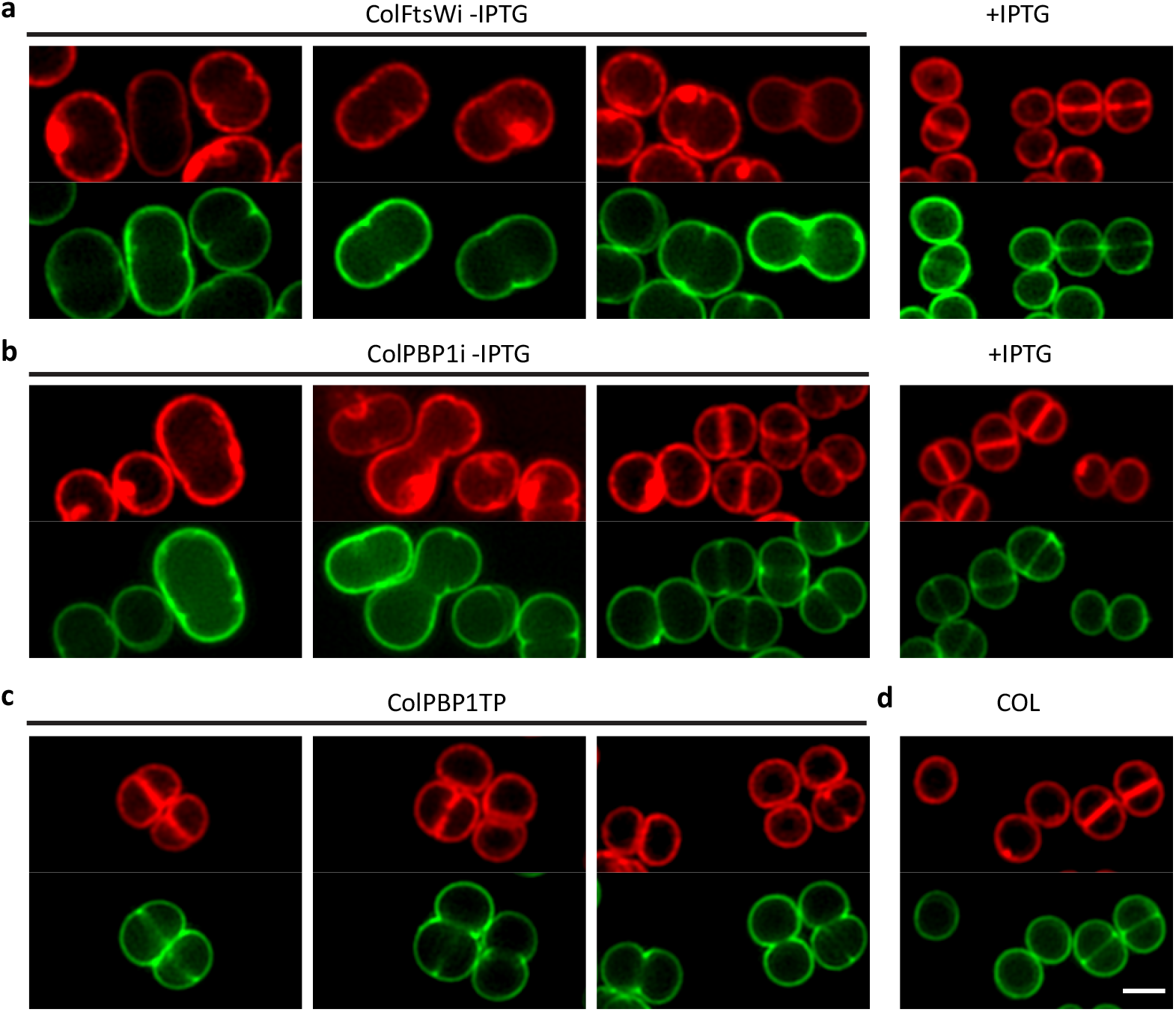
FtsW-PBP1 has a role in maintaining the near-spherical shape of cocci. a, b, SIM images of ColFtsWi (**a**) and ColPBP1i (**b**) grown in the absence and presence of IPTG, stained with Nile Red (membrane, Red) and Van-FL (cell wall, Green). **c, d,** SIM images of PBP1 transpeptidase mutant ColPBP1TP (**c**) and parental strain COL (**d**) stained with Nile Red (membrane, Red) and Van-FL (cell wall, Green). Scale bar, 1 µm.

Given that PBP1 TP activity is not required for growth, we tested whether the morphological phenotypes of the PBP1 depleted cells were dependent on PBP1 TP activity. Compared to the parental strain COL, ColPBP1TP cells showed cell separation defects, where daughter cells began to synthesise the future division septum while still attached (Fig. 3c,d, Supplementary Fig. 6c,d, white arrows). After completion of septum synthesis, cells did not quickly pop apart, as observed with COL, but instead slowly acquired an indentation at midcell. Importantly, the elongation phenotype, characteristic of the PBP1 depleted cells, was not observed in the PBP1TP mutant (Fig. 3c), suggesting that PBP1 has a second function besides its TP activity. A second line of evidence pointing to an early function of PBP1, separate from its role in PGN synthesis, comes from the fact that FtsW and PBP1 localise almost exclusively at midcell, but inward septum synthesis is not initiated until the lipid II flippase MurJ is recruited to the divisome^21^. Given the elongation phenotype of FtsW or PBP1 depleted cells, we reasoned that this cognate pair could have an initial structural role in divisome assembly and that cell elongation could arise from destabilisation of the divisome. We therefore determined the localisation of FtsZ and its negative regulator EzrA^23,24^ (as a proxy) upon depletion of FtsW and PBP1. Imaging of FtsZ^55-56^sGFP and EzrA-sGFP, in the absence of FtsW or PBP1, revealed the presence of multiple rings/arcs spread across the long axis of the cell, indicating that these proteins have a stabilising effect on the divisome, maintaining it at midcell (Fig. 4a, Supplementary Fig. 7, 8a,b). These extraneous Z-rings appeared capable of recruiting downstream proteins of the divisome, as multiple foci (representing multiple rings in 2D) were observed for FtsW, MurJ and RodA in the absence of PBP1 (Supplementary Fig. 7). Therefore it seemed likely that cell elongation was due to PGN incorporation organised by the mislocalised divisome. To test this, we observed the incorporation of FDAAs following depletion of either FtsW or PBP1. Initial imaging showed faint septal incorporation, with peripheral signal throughout the cell (Supplementary Fig. 9). Given that the low molecular weight PBP4 is responsible for the majority of peripheral FDAA incorporation in *S. aureus*^16^, it was possible that a signal corresponding to divisome localisation could have been masked by PBP4 activity. We therefore repeated these experiments in the absence of PBP4, and found that similarly to the divisome localisation, multiple PGN incorporation rings were visible along the long axis of the mutant (Fig. 5, Supplementary Fig. 8c). HADA staining of these strains expressing EzrA-sGFP confirmed that the insertion of PGN was directed by the mis-localised divisome, indicating that cell elongation is caused by lateral PGN insertion at defined regions of the cell (Fig. 4b, Supplementary Fig. 8d). Together these data support a model whereby both SEDS-bPBPs cognate pairs are responsible for maintaining the cell shape by directing peptidoglycan synthesis to the cell periphery or septum during the cell cycle of *S. aureus*.

**Figure 4.**
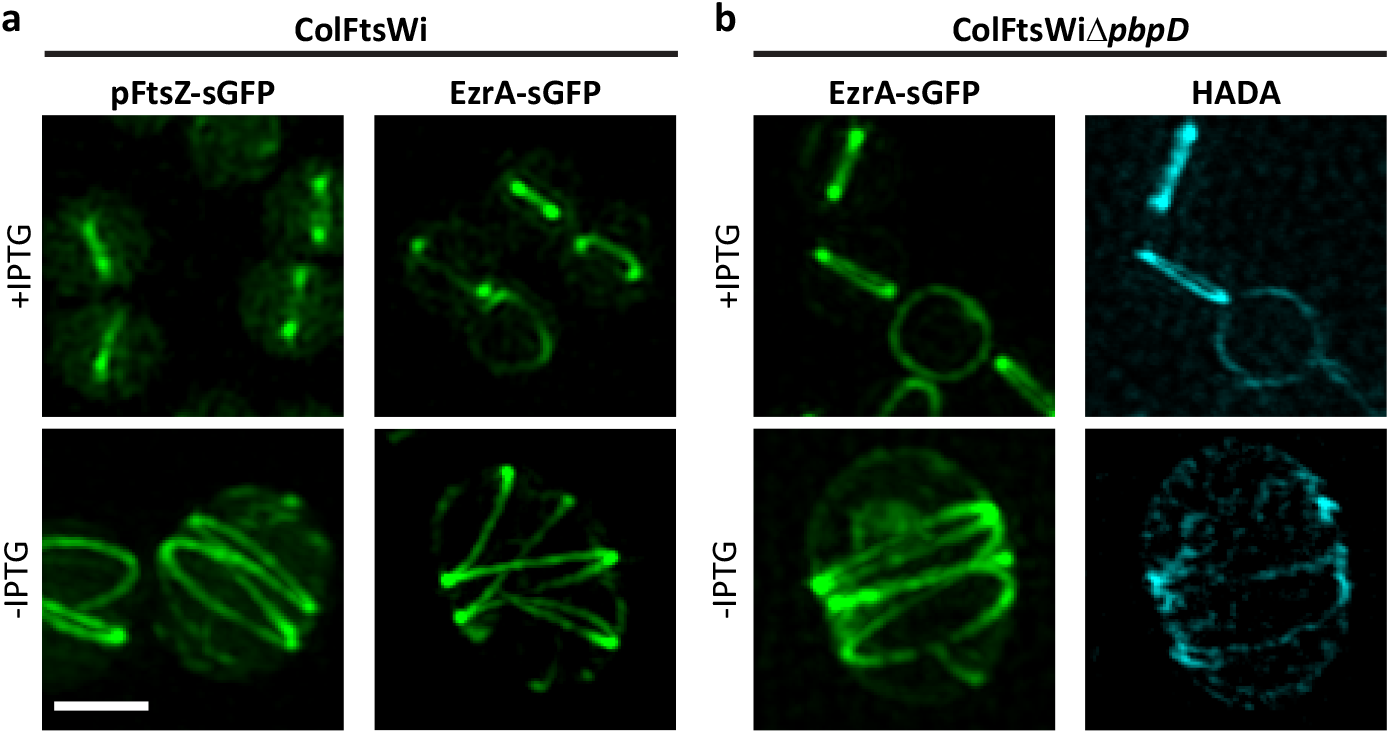
FtsW depletion leads to multiple divisome structures that incorporate PGN. a, FtsZ-sGFP and EzrA-sGFP localisation in ColFtsWipFtsZ-sGFP and ColFtsWiEzrA-sGFP cells depleted (-IPTG) or not (+IPTG) of FtsW. **b**, EzrA-sGFP (left) and PGN insertion visualised by HADA staining (right) in PBP4 null mutant ColFtsWiΔ*pbpD*EzrA-sGFP depleted (-IPTG) or not (+IPTG) of FtsW. Panels show a maximum intensity projection of a SIM Z-stack, except top images in panel a, that show a 2D-SIM image. Scale bar, 1 µm.

**Figure 5.**
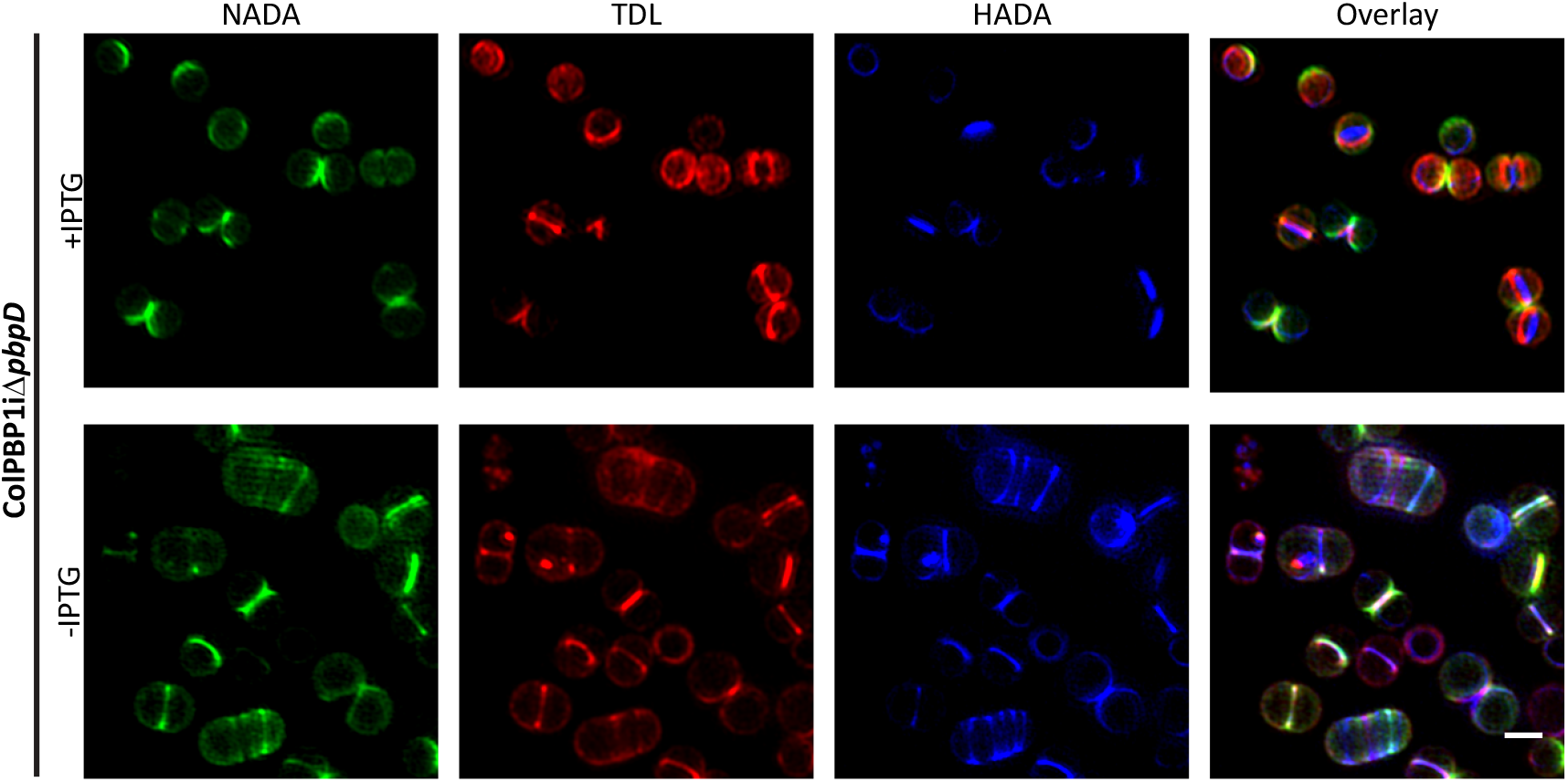
FtsW and PBP1 depleted cells elongate due to PGN incorporation at the lateral wall. SIM images of strain ColPBP1iΔ*pbpD* grown in the presence and absence of IPTG to deplete PBP1 and sequentially stained for 10 min (each) with fluorescent D-amino acids NADA, TDL and HADA, showing PGN incorporation in bands at the lateral wall. Scale bar, 1 µm.

## DISCUSSION

In rod shaped bacteria, the elongasome (via MreB) ensures synthesis along the long axis of the cell resulting in elongation, while the divisome (via FtsZ) promotes inward synthesis of the septum^7,25^. Each of these machineries includes a specific SEDS-bPBP pair required for lateral or septal synthesis of PGN^14^. Cocci like *S. aureus* lack a canonical elongasome including an MreB protein. However, they do encode two SEDS proteins whose role was so far unknown. We have now confirmed the existence of two SEDS-bPBP cognate pairs of interacting proteins. The RodA-PBP3 pair is recruited to midcell and promotes lateral insertion of PGN, leading to slight cell elongation, in a manner reminiscent of pre-septal PGN synthesis of *E. coli*^10^ or *Caulobacter crescentus*^26^ or peripheral PGN synthesis responsible for cell elongation in ovococci^8^, both of which are dependent on FtsZ. The second SEDS-bPBP pair, FtsW-PBP1, is essential for cell viability and is required for inward PGN synthesis at the division site. Interestingly, the essential function of PBP1 is not dependent on its transpeptidase activity, suggesting that this protein has a second function. While writing this manuscript, preprint data from the Walker laboratory demonstrated that *S. aureus* PBP1 stimulates the essential TG activity of FtsW, independently of its TP active site^12^, which could explain the essentiality of PBP1. This regulation of the activity of SEDS proteins and bPBPs may be essential to avoid the synthesis of uncrosslinked glycans^27^, which has been shown to lead to a toxic futile cycle of glycan synthesis and degradation in *E. coli*^28^. We now propose an additional mode of regulation of SEDS proteins by cognate bPBPs through protein localisation. Our data has shown that *S. aureus* PBP3 is required for the septal localisation of RodA, and most likely modulates its activity, as lack of the transpeptidase PBP3 causes a similar increase in susceptibility to moenomycin (an inhibitor of classical transglycosylases) as the lack of RodA. We also found that FtsW-PBP1 pair has an additional function in stabilising the divisome. FtsW has been previously suggested to have a role in divisome activation in *E. coli* and *C. crescentus*^29,30^. Here we showed that *S. aureus* FtsW-PBP1 pair is required to maintain the divisome at midcell, as depletion of these proteins led to multiple rings/arcs of FtsZ or EzrA present across the long axis of the cell. FtsW and PBP1 arrive early to the divisome of *S. aureus*, even before the lipid II flippase MurJ^21^. It is therefore possible that a major function of FtsW-PBP1 is to stabilise the early divisome before this SEDS-bPBP pair initiates TG and TP activity in the orientation perpendicular to the membrane required to initiate septum synthesis. In the absence of these proteins, the normally compact midcell divisome mislocalises to the peripheral wall, leading to cell elongation.

In conclusion, we propose that *S. aureus* maintains its slightly ellipsoid shape through the coordinated activities of two cognate SEDS-bPBP pairs that insert new PGN in two distinct fashions at the midcell. Newborn cells incorporate PGN around the entire surface^16,31^. Just before septum inward growth is initiated, RodA-PBP3 localises at midcell and incorporates PGN at the lateral wall, while FtsW-PBP1 stabilises assembly of the divisome during initial stages of cytokinesis. Importantly, the transpeptidase activity of PBP1 is required to ensure that cell wall maturation of the closed septum concludes with rapid daughter cell separation and reshaping of the septum. Together these SEDS-bPBP pairs localise PGN incorporation and this balance between lateral and septal PGN synthesis is crucial in conserving the *S. aureus* cell shape. Our findings suggest that cell shape development in cocci, ovococci and rods presents more similarities than previously thought, with the three morphologies requiring protein complexes for elongation and division.

## ACKNOWLEDGEMENTS

We thank Leonilde Moreira (IST) for hosting laboratory work at Instituto Superior Técnico, University of Lisbon, Lisbon, Portugal, Sérgio R. Filipe (FCT-NOVA) for helpful discussions, Sara Bonucci and Erin M. Tranfield (Electron Microscopy Facility, IGC) for technical expertise and sample processing and André Bernardo, Inês Jorge, Dominika Kadziolka and Kinga Witana for help in the construction of some plasmids. This study was funded by the European Research Council through grant ERC-2017-CoG-771709 (to MGP), Project LISBOA-01-0145FEDER-007660 Microbiologia Molecular, Estrutural e Celular (to ITQB-NOVA), the Portuguese Platform of Bioimaging PPBI-POCI-01-0145-FEDER-022122, Researcher Contract No. IF/00386/2015 (to FF) and FCT fellowships SFRH/BPD/95031/2013 (NTR) and SFRH/BD/52204/2013 (ACT). WGS analysis at the Genomics Unit of Instituto Gulbenkian de Ciencia (Oeiras, Portugal) was supported by ONEIDA project (LISBOA-01-0145-FEDER-016417) co-funded by FEEI - “Fundos Europeus Estruturais e de Investimento” from “Programa Operacional Regional Lisboa 2020” and by national funds from FCT - “Fundação para a Ciência e a Tecnologia.”

## AUTHOR CONTRIBUTIONS

N.T.R., A.C.T. and M.G.P. designed the research. N.T.R. and A.C.T. performed all experiments with the exception of FLIM data acquisition and analysis which was performed by F.F., and HPLC muropeptide analysis performed by A.J. B.M.S. developed software for image analysis of seFRET data. N.T.R., A.C.T. and P.R. constructed strains. A.J. and P.R. analysed WGS data. J.M.M and A.R.P performed preliminary experiments and analysed data. R.G.S. performed preliminary experiments. M.S.V. contributed new reagents (FDAAs). N.T.R., A.C.T. and M.G.P. analysed the overall data. B.M.S. analysed seFRET data quantified by in-house developed software. N.T.R., A.C.T. and M.G.P. wrote the manuscript.

---

**Supplementary Figure 1.**
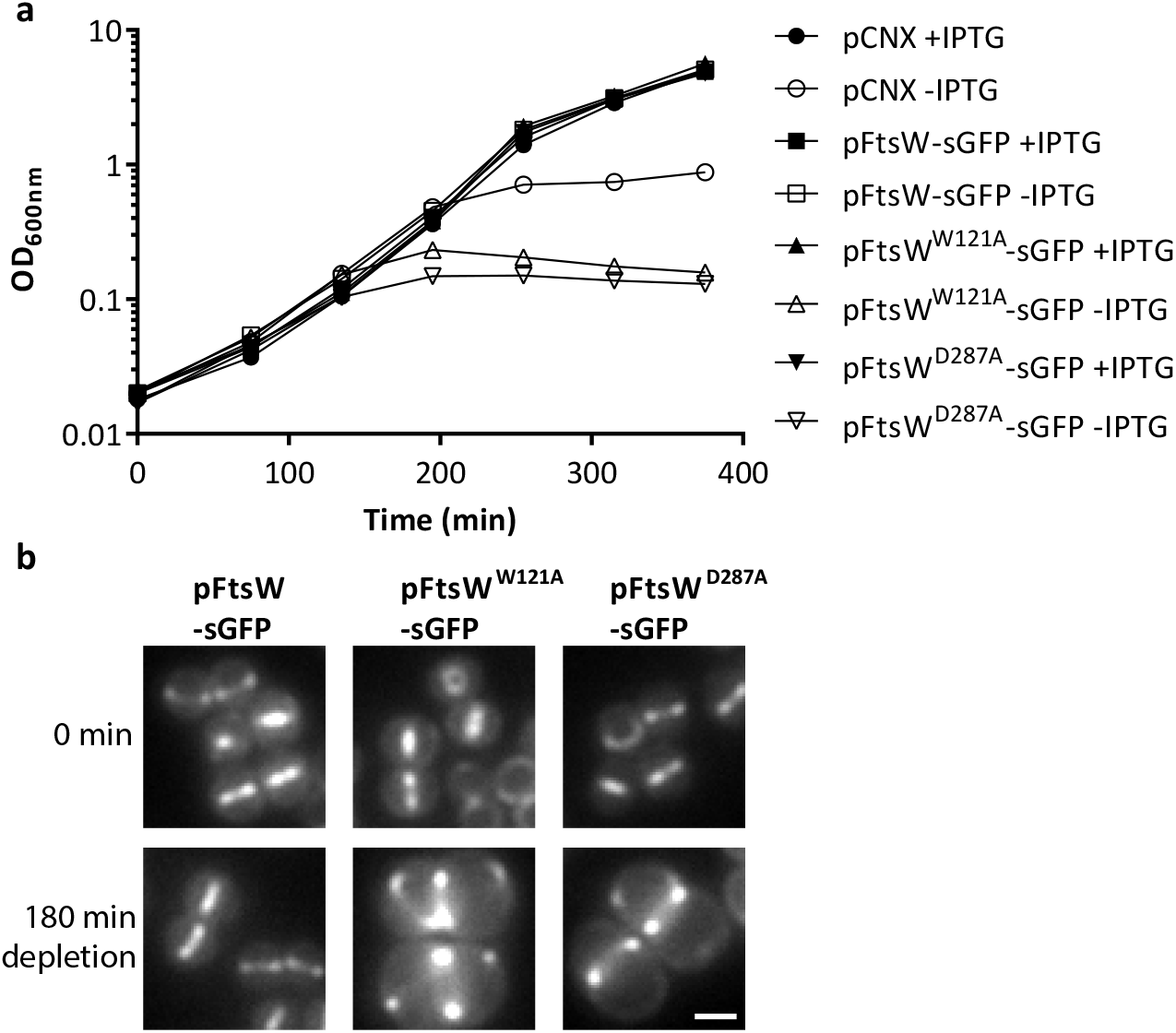
FtsW^W121A^ and FtsW^D287A^ are non-functional. a, Growth curves of strain ColFtsWi encoding a chromosomal copy of FtsW under the control of an IPTG inducible promoter and complemented with the empty vector pCNX, the same vector encoding a sGFP derivative of native FtsW (pCNX-ftsWsgfp) or two active site mutants (pCNX-ftsW^W121A^sgfp and pCNX-ftsW^D287A^sgfp). Strains were grown in the presence or absence of IPTG. **b**, Epi-fluorescence microscopy images of ColFtsWi complemented with FtsW-sGFP derivatives expressed from a vector (pCNX-ftsWsgfp, pCNX-ftsW^W121A^sgfp and pCNX-ftsW^D287A^sgfp). Images were taken before (0 min) and after (180 min) depletion of chromosomally encoded FtsW. Scale bar, 1 µm.

**Supplementary Figure 2.**
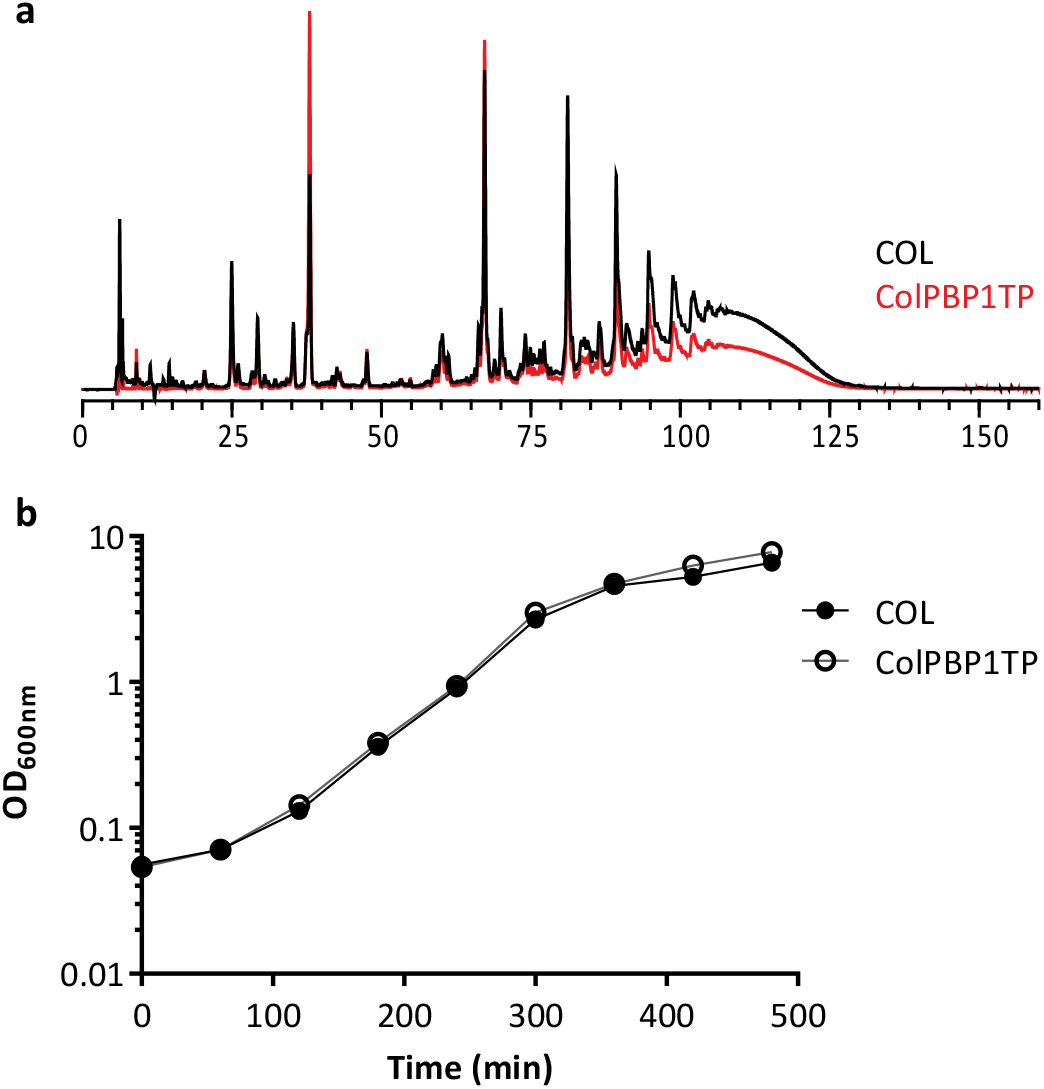
PBP1 TP mutant shows reduced PGN crosslinking while maintaining normal growth rate. a, HPLC analysis of peptidoglycan muropeptides from parental strain COL (black) and PBP1 TP mutant ColPBP1TP (red). **b**, Growth curve of COL and ColPBP1TP in TSB medium.

**Supplementary Figure 3.**
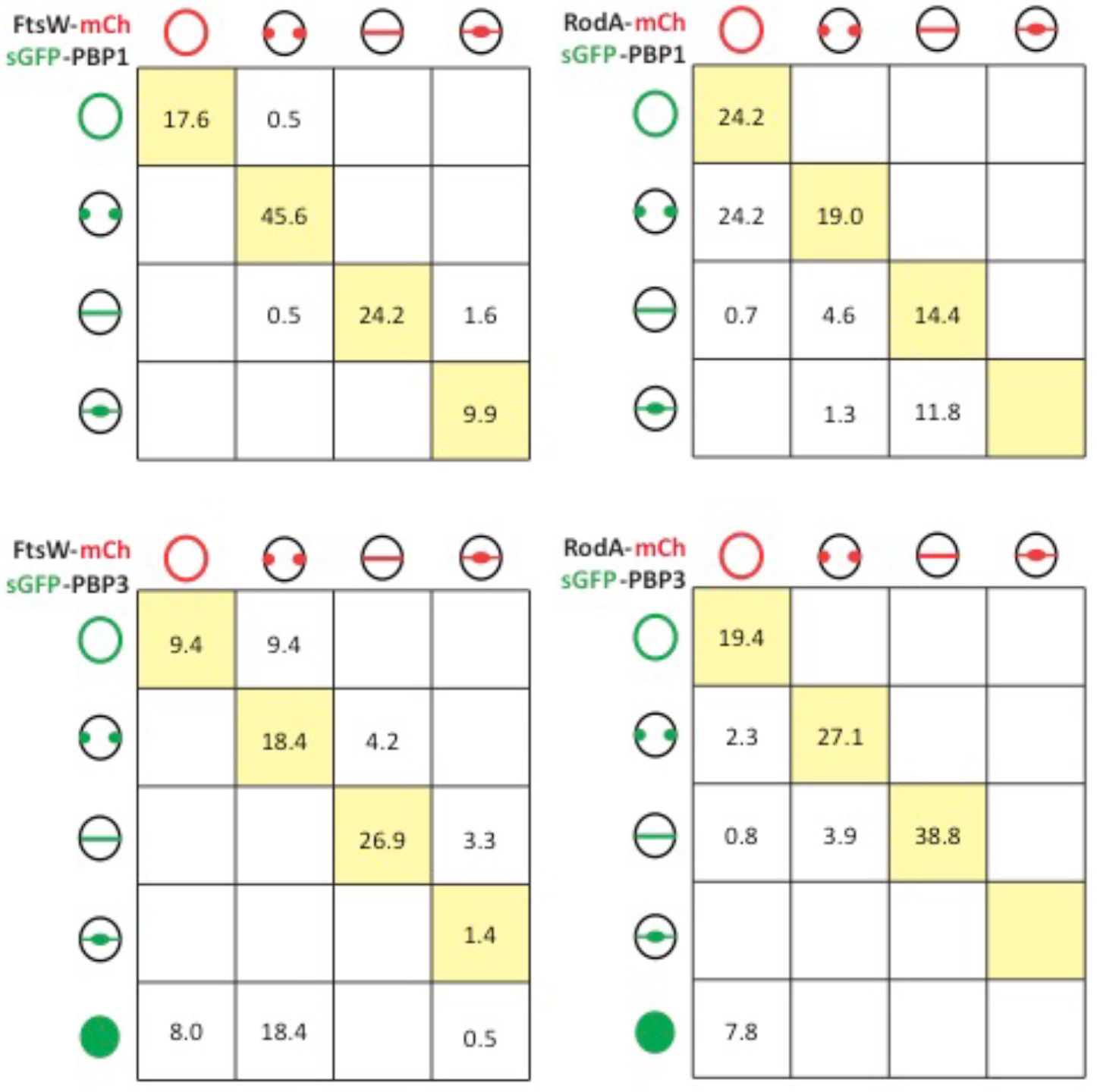
FtsW-mCherry/GFP-PBP1 and RodA-mCherry/sGFP-PBP3 colocalise in the majority of cells. Tables correlate the localisation of mCherry derivatives of SEDS proteins FtsW and RodA with sGFP derivatives of bPBPs PBP1 and PBP3, in strains ColWP1, ColWP3, ColP1pA and ColP3pA. Yellow squares show the percentage of cells with colocalisation of the SEDS and bPBP proteins at different stages of the cell cycle. Data suggests that PBP1 arrives to the septum before RodA (top right table) and that FtsW arrives to the septum before PBP3 (bottom left corner). *n* >100 cells for each strain, in each of three biological repeats.

**Supplementary Figure 4.**
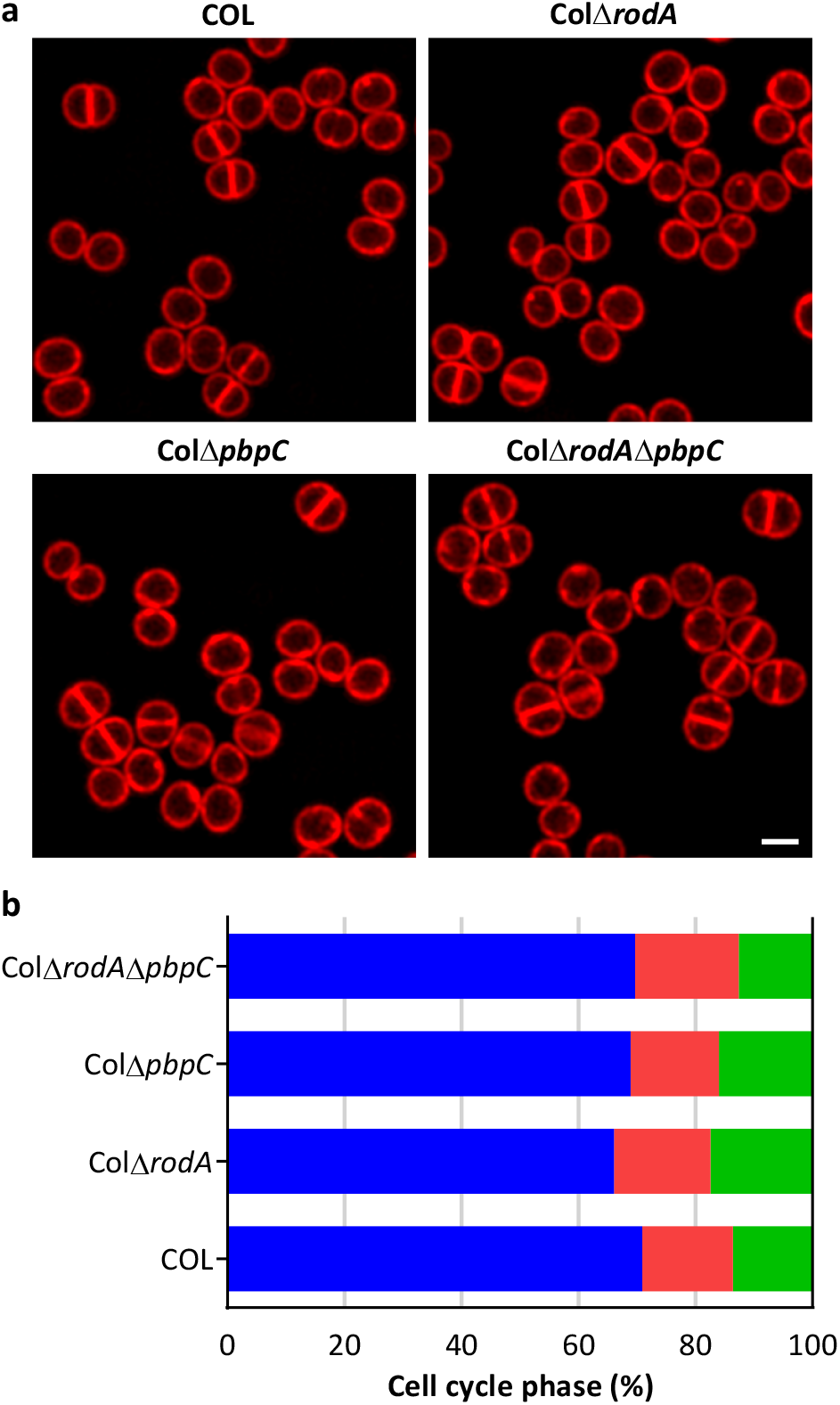
Absence of RodA and PBP3 has no effect on cell cycle progression. a, SIM images of mid-exponential phase cells of COL, ColΔ*rodA*, ColΔ*pbpC* and ColΔ*rodA*Δ*pbpC* stained with Nile Red. Scale bar, 1µm. **b**, Percentage of cells in cell cycle phase 1 (cells with no septum, blue), phase 2 (cells undergoing septum synthesis, red) and phase 3 (cells with a complete septum, green), determined from SIM images of Nile Red stained cells. Three independent experiments were performed and for each experiment at least three different images were analysed (*n*=100 for each image, *n*>300 for each experiment). The average of the three experiments is shown in the graph and corresponds to the fraction of the cell cycle spent in each phase.

**Supplementary Figure 5.**
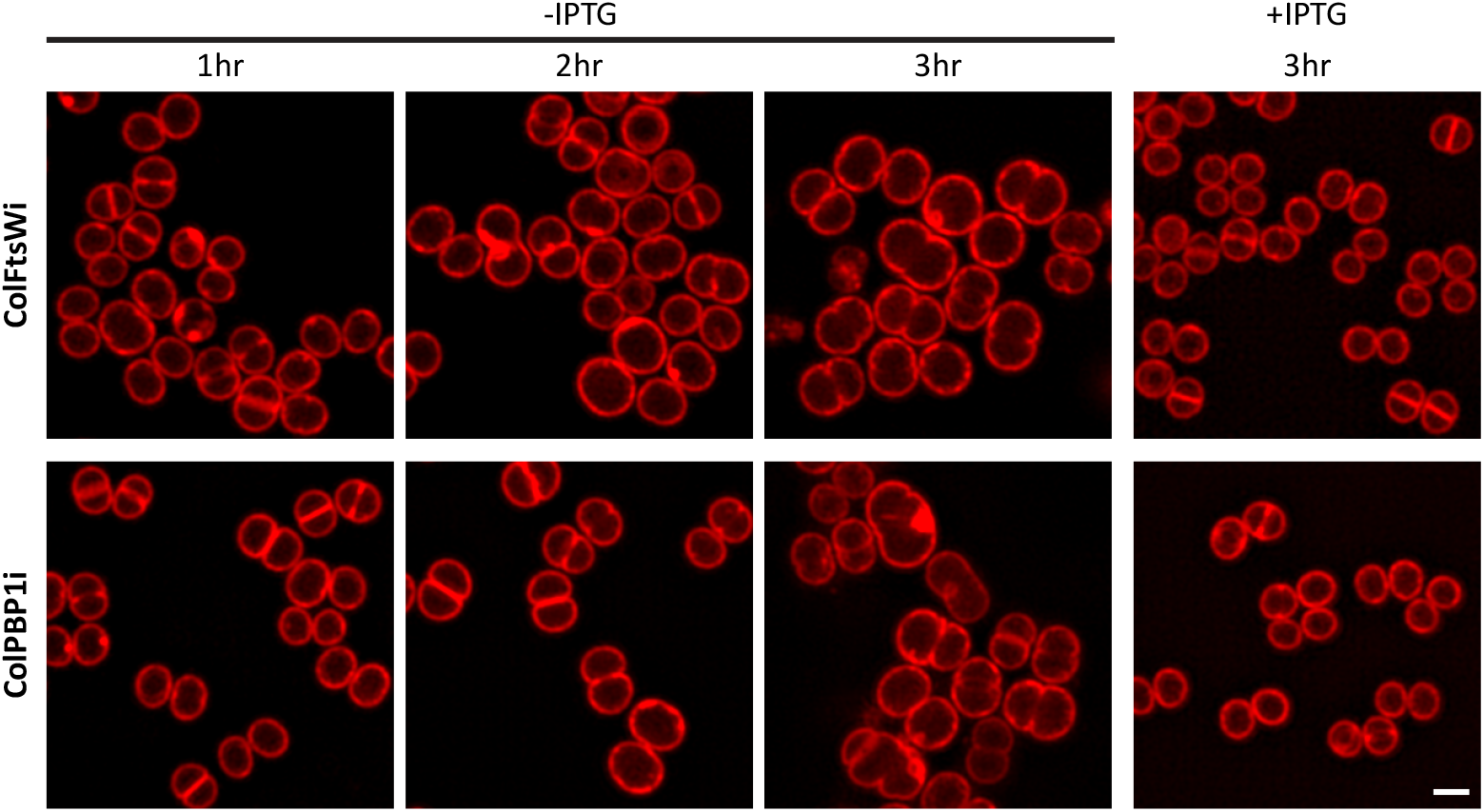
Depletion of FtsW and PBP1 leads to cell elongation. SIM images following 1 h, 2 h and 3 h of FtsW (ColFtsWi) or PBP1 (ColPBP1i) depletion (-IPTG). Strains grown in the presence of IPTG were imaged at 3 hr. Scale bar, 1 µm.

**Supplementary Figure 6.**
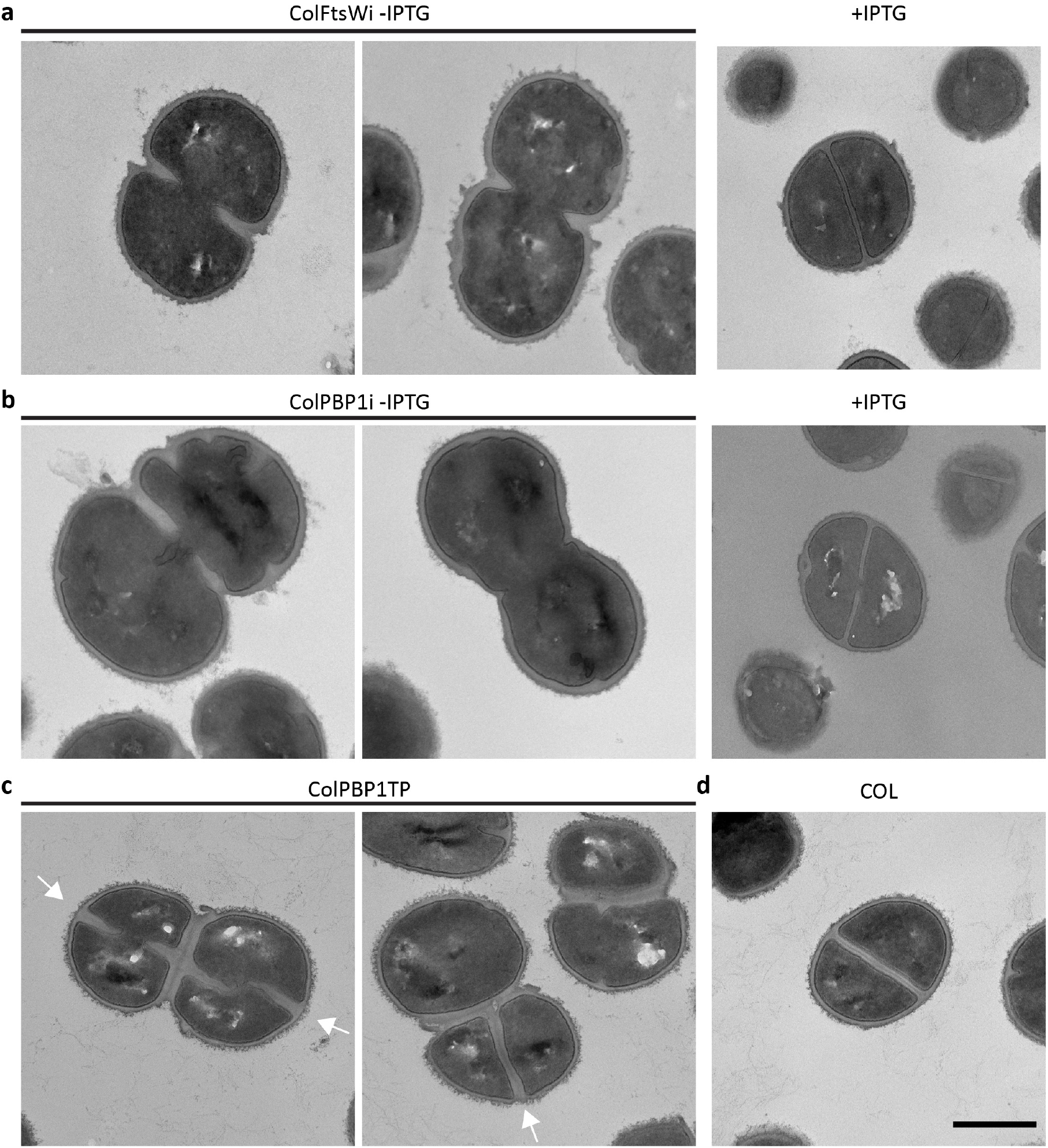
Depletion of PBP1 or FtsW and mutation of TP active site of PBP1 leads to cell division defects. a, b, Transmission Electron Microscopy (TEM) images of ColFtsWi (**a**) and ColPBP1i (**b**) grown in the absence and presence of IPTG. **c, d** TEM images of PBP1 TP mutant ColPBP1TP (**c**) and parental strain COL (**d**). Arrows in (**c**) indicate division septa present prior to daughter cell separation. Scale bar, 0.5 µm.

**Supplementary Figure 7.**
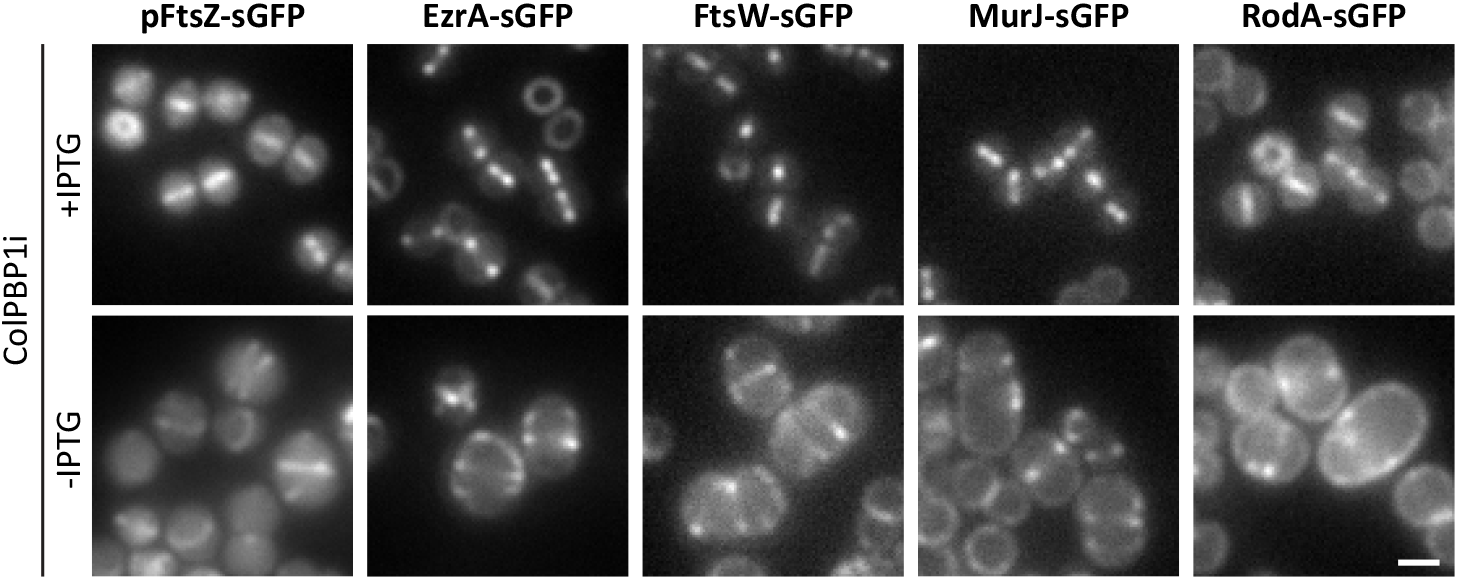
PBP1 depletion leads to mislocalisation of divisome proteins. Epi-fluorescence microscopy images of (left to right) ColPBP1ipFtsZ-sGFP, ColPBP1iEzrA-sGFP, ColPBP1iFtsW-sGFP, ColPBP1iMurJ-sGFP and ColPBP1iRodA-sGFP grown in the presence (+IPTG) and absence of IPTG for PBP1 depletion (-IPTG). The multiple foci observed in the absence of IPTG likely correspond to multiple rings. Scale bar, 1 µm.

**Supplementary Figure 8.**
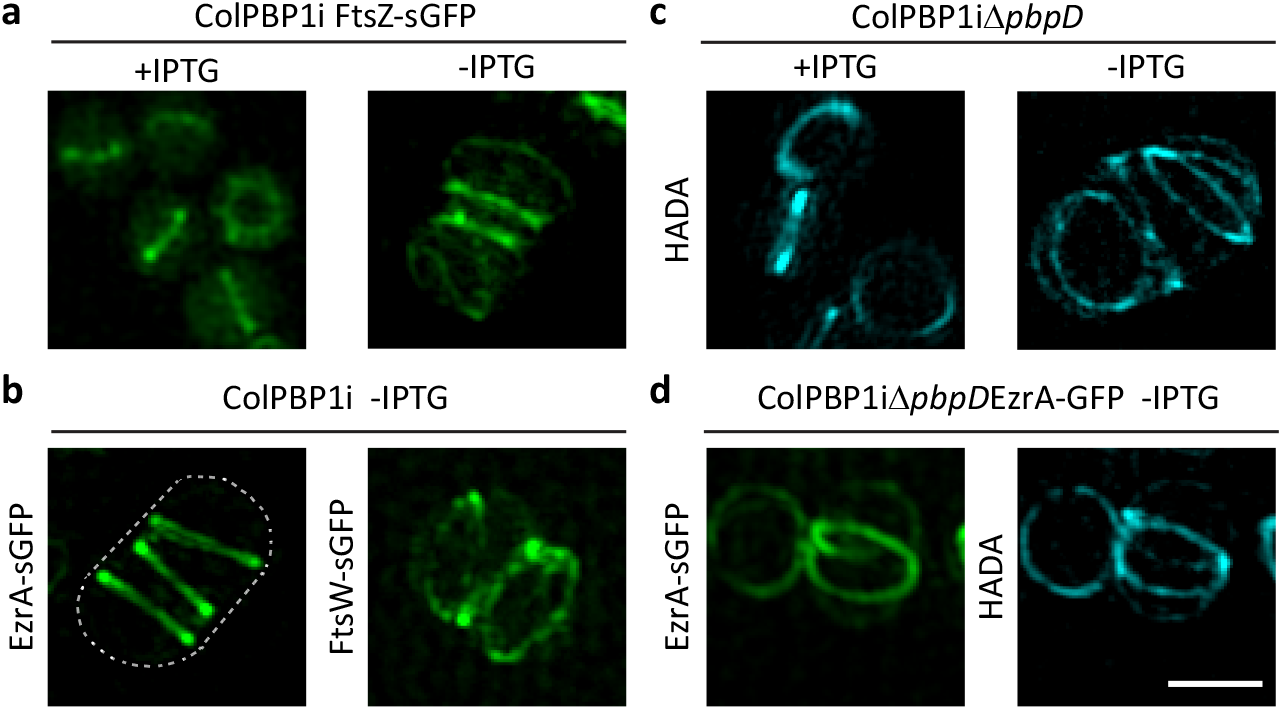
PBP1 depletion leads to multiple divisome structures that incorporate PGN. a, FtsZ-sGFP localisation in ColPBP1ipFtsZ-sGFP strain depleted (-IPTG) or not (+IPTG) of PBP1. **b**, EzrA-sGFP and FtsW-sGFP localisation in ColPBP1iEzrA-sGFP and ColPBP1iFtsW-sGFP strains depleted of PBP1. Contour of the cell is indicated by a dashed line. **c**, PGN insertion visualised by HADA staining in PBP4 null mutant strain ColPBP1iΔ*pbpD* depleted (-IPTG) or not (+IPTG) of PBP1. **d**, Co-localisation of EzrA-sGFP (left) and PGN insertion visualised by HADA staining (right) in ColPBP1iΔ*pbpD*EzrA-sGFP depleted of PBP1. Left panel of (a) and (c) are 2D-SIM images while the other panels are a maximum intensity projection of a SIM Z-stack. Scale bar, 1 µm.

**Supplementary Figure 9.**
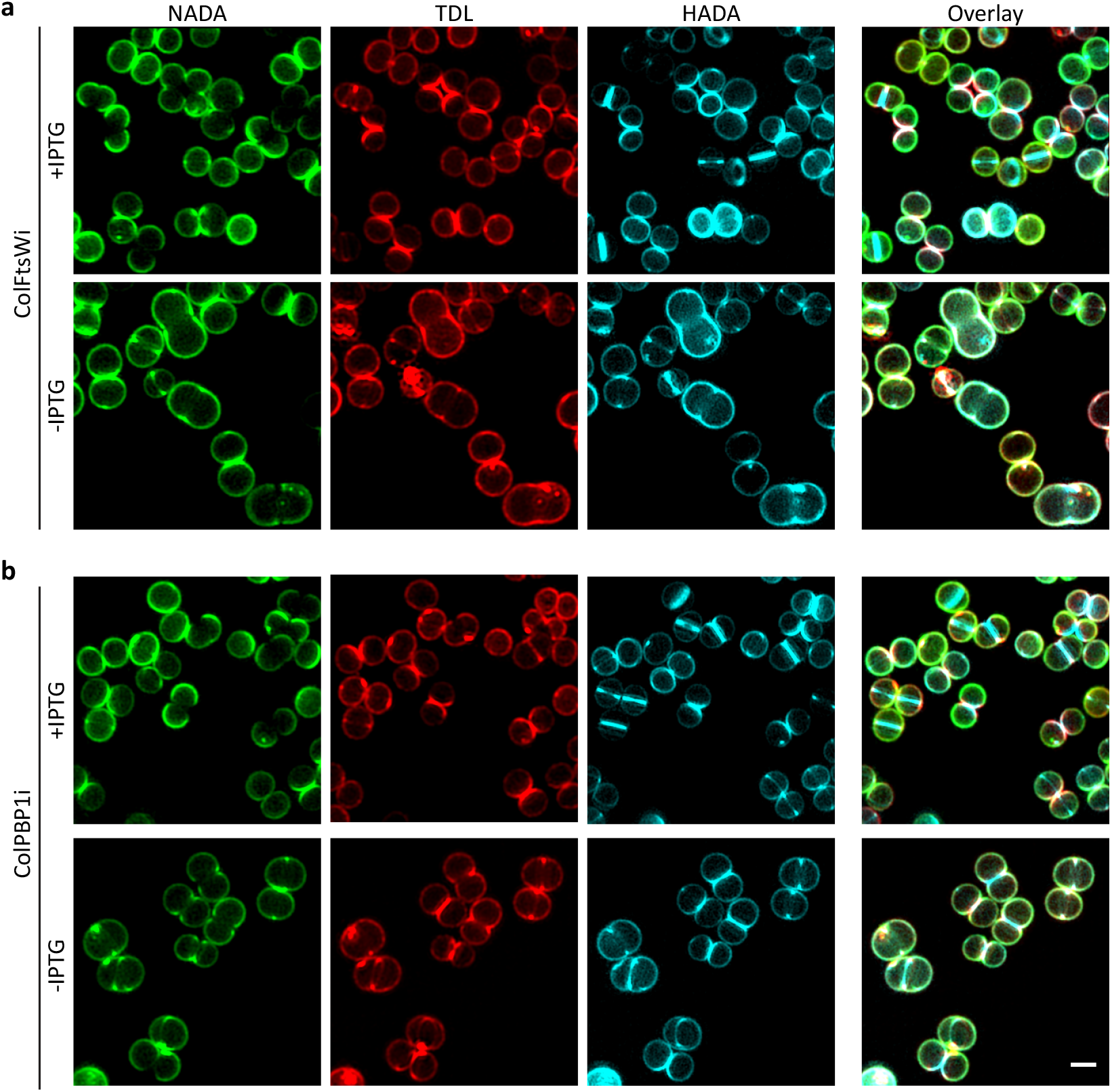
PGN incorporation over time in cells depleted of FtsW or PBP1. a, b, SIM images of strains ColFtsWi (a) and ColPBP1i (b) grown in the presence and absence of IPTG and sequentially stained for 10 min (each) with fluorescent D-amino acids NADA, TDL and HADA. Scale bar, 1 µm.

## BIBLIOGRAPHY

1. Egan, A. J., Cleverley, R. M., Peters, K., Lewis, R. J. & Vollmer, W. Regulation of bacterial cell wall growth. FEBS J 284, 851-867, doi:10.1111/febs.13959 (2017).

2. Chastanet, A. & Carballido-Lopez, R. The actin-like MreB proteins in Bacillus subtilis: a new turn. Front Biosci (Schol Ed) 4, 1582–1606 (2012).

3. Domínguez-Escobar, J. et al. Processive movement of MreB-associated cell wall biosynthetic complexes in bacteria. Science 333, 225-228, doi:10.1126/science.1203466 (2011).

4. Garner, E. C. et al. Coupled, circumferential motions of the cell wall synthesis machinery and MreB filaments in *B. subtilis*. Science 333, 222-225, doi:10.1126/science.1203285 (2011).

5. van Teeffelen, S. et al. The bacterial actin MreB rotates, and rotation depends on cell-wall assembly. Proc Natl Acad Sci U S A 108, 15822-15827, doi:10.1073/pnas.1108999108 (2011).

6. den Blaauwen, T., Hamoen, L. W. & Levin, P. A. The divisome at 25: the road ahead. Curr Opin Microbiol 36, 85-94, doi:10.1016/j.mib.2017.01.007 (2017).

7. Egan, A. J. & Vollmer, W. The physiology of bacterial cell division. Annals of the New York Academy of Sciences 1277, 8-28, doi:10.1111/j.1749-6632.2012.06818.x (2013).

8. Massidda, O., Novakova, L. & Vollmer, W. From models to pathogens: how much have we learned about *Streptococcus pneumoniae* cell division? Environ Microbiol 15, 3133-3157, doi:10.1111/1462-2920.12189 (2013).

9. Pinho, M. G., Kjos, M. & Veening, J.-W. How to get (a)round: mechanisms controlling growth and division of coccoid bacteria. Nat. Rev. Microbiol. 11, 601-614, doi:10.1038/nrmicro3088 (2013).

10. Typas, A., Banzhaf, M., Gross, C. A. & Vollmer, W. From the regulation of peptidoglycan synthesis to bacterial growth and morphology. Nat Rev Microbiol 10, 123-136, doi:10.1038/nrmicro2677 (2011).

11. Meeske, A. J. et al. SEDS proteins are a widespread family of bacterial cell wall polymerases. Nature 537, 634–638, doi:10.1038/nature19331 (2016).

12. Taguchi, A. et al. FtsW is a peptidoglycan polymerase that is activated by its cognate penicillin-binding protein. Preprint at https://www.biorxiv.org/content/early/2018/06/29/358663 (2018).

13. Emami, K. et al. RodA as the missing glycosyltransferase in *Bacillus subtilis* and antibiotic discovery for the peptidoglycan polymerase pathway. Nat Microbiol 2, 16253, doi:10.1038/nmicrobiol.2016.253 (2017).

14. Cho, H. et al. Bacterial cell wall biogenesis is mediated by SEDS and PBP polymerase families functioning semi-autonomously. Nat Microbiol 1, 16172, doi:10.1038/nmicrobiol.2016.172 (2016).

15. Scheffers, D. J. & Pinho, M. G. Bacterial cell wall synthesis: New insights from localization studies. Microbiol. Mol. Biol. Rev. 69, 585-607, doi:10.1128/MMBR.69.4.585-607.2005 (2005).

16. Monteiro, J. M. et al. Cell shape dynamics during the staphylococcal cell cycle. Nat Commun 6, 8055, doi:10.1038/ncomms9055 (2015).

17. Pereira, S. F., Henriques, A. O., Pinho, M. G., de Lencastre, H. & Tomasz, A. Role of PBP1 in cell division of *Staphylococcus aureus*. J. Bacteriol. 189, 3525-3531, doi:10.1128/JB.00044-07 (2007).

18. Pinho, M. G., de Lencastre, H. & Tomasz, A. Cloning, characterization, and inactivation of the gene *pbpC*, encoding penicillin-binding protein 3 of *Staphylococcus aureus*. J Bacteriol 182, 1074-1079, doi:10.1128/JB.182.4.1074-1079.2000 (2000).

19. Pereira, S. F. F., Henriques, A. O., Pinho, M. G., de Lencastre, H. & Tomasz, A. Evidence for a dual role of PBP1 in the cell division and cell separation of *Staphylococcus aureus*. Mol Microbiol 72, 895-904, doi:10.1111/j.1365-2958.2009.06687.x (2009).

20. Sassine, J. et al. Functional redundancy of division specific penicillin-binding proteins in *Bacillus subtilis*. Mol Microbiol 106, 304-318, doi:10.1111/mmi.13765 (2017).

21. Monteiro, J. M. et al. Peptidoglycan synthesis drives an FtsZ-treadmilling-independent step of cytokinesis. Nature 554, 528-532, doi:10.1038/nature25506 (2018).

22. Kuru, E. et al. In situ Probing of Newly Synthesized Peptidoglycan in Live Bacteria with Fluorescent D-Amino Acids. Angew. Chem. Int Ed. Engl. 51, 12519-12523, doi:10.1002/anie.201206749 (2012).

23. Levin, P. A., Kurtser, I. G. & Grossman, A. D. Identification and characterization of a negative regulator of FtsZ ring formation in *Bacillus subtilis*. Proc Natl Acad Sci U S A 96, 9642–9647 (1999).

24. Jorge, A. M., Hoiczyk, E., Gomes, J. P. & Pinho, M. G. EzrA Contributes to the Regulation of Cell Size in *Staphylococcus aureus*. PLoS ONE 6, e27542, doi:10.1371/journal.pone.0027542 (2011).

25. den Blaauwen, T., de Pedro, M. A., Nguyen-Disteche, M. & Ayala, J. A. Morphogenesis of rod-shaped sacculi. FEMS Microbiol Rev 32, 321-344, doi:10.1111/j.1574-6976.2007.00090.x (2008).

26. Aaron, M. et al. The tubulin homologue FtsZ contributes to cell elongation by guiding cell wall precursor synthesis in *Caulobacter crescentus*. Mol Microbiol 64, 938-952, doi:10.1111/j.1365-2958.2007.05720.x (2007).

27. Rohs, P. D. A. et al. An activation pathway governs cell wall polymerization by a bacterial morphogenic machine. Preprint at https://www.biorxiv.org/content/early/2018/06/29/359208 (2018).

28. Cho, H., Uehara, T. & Bernhardt, T. G. Beta-lactam antibiotics induce a lethal malfunctioning of the bacterial cell wall synthesis machinery. Cell 159, 1300-1311, doi:10.1016/j.cell.2014.11.017 (2014).

29. Du, S., Pichoff, S. & Lutkenhaus, J. FtsEX acts on FtsA to regulate divisome assembly and activity. Proc Natl Acad Sci U S A 113, E5052-5061, doi:10.1073/pnas.1606656113 (2016).

30. Modell, J. W., Kambara, T. K., Perchuk, B. S. & Laub, M. T. A DNA damage-induced, SOS-independent checkpoint regulates cell division in *Caulobacter crescentus*. PLoS Biol 12, e1001977, doi:10.1371/journal.pbio.1001977 (2014).

31. Lund, V. A. et al. Molecular coordination of *Staphylococcus aureus* cell division. eLife 7, doi:10.7554/eLife.32057 (2018).

