## Supplementary Tables for "SEDS-bPBP pairs direct Lateral and Septal Peptidoglycan Synthesis in *Staphylococcus aureus*"

### Supplementary Table 1 – Strains and plasmids used in this study

| **Strains and plasmids** | **Description** | **Source or reference** |
|---|---|---|
| ***Escherichia coli*** | | |
| DC10B | Δ*dcm* in DH10B background; Dam methylation only; for cloning | 1 |
| ***Staphylococcus aureus*** | | |
| RN4220 | restriction-negative derivative of NCTC8325-4 | 2 |
| COL | HA-MRSA | 3 |
| ColΔ*rodA* | *rodA* deletion of parental strain COL | This work |
| ColΔ*pbpC* | *pbpC* deletion of parental strain COL | This work |
| ColΔ*rodA*Δ*pbpC* | *rodA* and *pbpC* deletions of parental strain COL | This work |
| ColpMGPII | COL with pMGPII; $Cam^r$ | |
| ColFtsWhigh | COL *spa*::$P_{spac}$*ftsW* | This work |
| ColFtsWlow | COL Δ*ftsW* *spa*::$P_{spac}$*ftsW* | This work |
| ColFtsWi | COL Δ*ftsW* *spa*::$P_{spac}$*ftsW* with pMGPII; $Cam^r$ | This work |

| **Strains and plasmids** | **Description** | **Source or reference** |
|---|---|---|
| ColFtsWiΔ*pbpD* | COL Δ*ftsW* Δ*pbpD spa::*$P_{spac}$*ftsW* with pMGPII; $Cam^r$ | This work |
| ColPBP1high | COL *spa::*$P_{spac}$*pbpA* | This work |
| ColPBP1low | COL Δ*pbpA spa::*$P_{spac}$*pbpA* | This work |
| ColPBP1i | COL Δ*pbpA spa::*$P_{spac}$*pbpA* with pMGPII; $Cam^r$ | This work |
| ColPBP1iΔ*pbpD* | COL Δ*pbpA* Δ*pbpD spa::*$P_{spac}$*pbpA* with pMGPII; $Cam^r$ | This work |
| ColPBP1TP | COL *pbpA*::*pbpA*$^{S314A}$ | This work |
| ColFtsW-mCherry | COL *ftsW::ftsW-mCherry* | 4 |
| ColFtsW-sGFP | COL *ftsW::ftsW-sgfp* | 4 |
| ColsGFP-PBP1 | COL *pbpA*::*sgfp-pbpA* | 4 |
| ColsGFP-PBP3 | COL *pbpC*::*sgfp-pbpC* | 4 |
| ColMurJ-mCherry | COL *murJ::murJ-mCherry* | 4 |
| ColRodA-sGFP | COL *rodA*::*rodA-sgfp* | 4 |
| ColRodA-mCherry | COL *rodA*::*rodA-mCherry* | This work |
| ColpCNX | COL with pCNX; $Kan^r$ | This work |
| ColpFtsW-mCherry | COL with pCNX-ftsWmCh; $Kan^r$ | This work |

| Strains and plasmids | Description | Source or reference |
|---|---|---|
| ColpMurJ-mCherry | COL with pCNX-murJmCh; $Kan^r$ | |
| ColpsGFP-PBP1 | COL with pCNX-sgfppbpA; $Kan^r$ | This work |
| ColpsGFP-PBP3 | COL with pCNX-sgfppbpC; $Kan^r$ | This work |
| ColpRodA-mCherry | COL with pCNX-rodAmCh; $Kan^r$ | This work |
| ColpmCherry-sGFP | COL with pCNX-mChsgfp; $Kan^r$ | This work |
| ColpmCherry-sGFP-TM | COL with pCNX-mChsgfpTM; $Kan^r$ | This work |
| ColWP1 | COL *ftsW::ftsW-mCherry pbpA::sgfp-pbpA* | This work |
| ColWP3 | COL *ftsW::ftsW-mCherry pbpC::sgfp-pbpC* | This work |
| ColP1pA | COL *pbpA::sgfp-pbpA* pCNX-rodAmCh | This work |
| ColP3pA | COL *pbpC::sgfp-pbpC* pCNX-rodAmCh | This work |
| ColpWP1 | COL with pCNX-WP1; $Kan^r$ | This work |
| ColpJP1 | COL with pCNX-JP1; $Kan^r$ | This work |
| ColpAP1 | COL with pCNX-AP1; $Kan^r$ | This work |
| ColpAP3 | COL with pCNX-AP3; $Kan^r$ | This work |
| ColFtsWiEzrA-sGFP | COL Δ*ftsW spa::*$P_{spac}$*ftsW ezrA::ezrA-sgfp* with pMGPII; $Cam^r$ | This work |

| Strains and plasmids | Description | Source or reference |
| --- | --- | --- |
| ColFtsWiΔ*pbpD*EzrA-sGFP | COL Δ*ftsW* Δ*pbpD spa::*$P_{spac}$*ftsW ezrA::ezrA-sgfp* with pMGPII; $Cam^r$ | This work |
| ColFtsWipCNX | COL Δ*ftsW spa::*$P_{spac}$*ftsW* with pMGPII and pCNX; $Cam^r$, $Kan^r$ | This work |
| ColFtsWipFtsW-sGFP | COL Δ*ftsW spa::*$P_{spac}$*ftsW* with pMGPII and pCNX-ftsWsgfp; $Cam^r$, $Kan^r$ | This work |
| ColFtsWipFtsW$^{W121A}$-sGFP | COL Δ*ftsW spa::*$P_{spac}$*ftsW* with pMGPII and pCNX-ftsW$^{W121A}$sgfp; $Cam^r$, $Kan^r$ | This work |
| ColFtsWipFtsW$^{D287A}$-sGFP | COL Δ*ftsW spa::*$P_{spac}$*ftsW* with pMGPII and pCNX-ftsW$^{D287A}$sgfp; $Cam^r$, $Kan^r$ | This work |
| ColFtsWipFtsZ-sGFP | COL Δ*ftsW spa::*$P_{spac}$*ftsW* with pMGPII with pCN-ftsZ$^{55-56}$sGFP; $Cam^r$, $Kan^r$ | This work |
| ColPBP1iEzrA-sGFP | COL Δ*pbpA spa::*$P_{spac}$*pbpA ezrA::ezrA-sgfp* with pMGPII; $Cam^r$ | This work |
| ColPBP1iΔ*pbpD*EzrA-sGFP | COL Δ*pbpA* Δ*pbpD spa::*$P_{spac}$*pbpA ezrA::ezrA-sgfp* with pMGPII; $Cam^r$ | This work |
| ColPBP1iFtsW-sGFP | COL Δ*pbpA spa::*$P_{spac}$*pbpA ftsW::ftsW-sgfp* with pMGPII; $Cam^r$ | This work |
| ColPBP1ipFtsZ-sGFP | COL Δ*pbpA spa::*$P_{spac}$*pbpA* with pMGPII and pCN-ftsZ$^{55-56}$sGFP; $Cam^r$, $Kan^r$ | This work |
| ColPBP1iMurJ-sGFP | COL Δ*pbpA spa::*$P_{spac}$*pbpA murJ::murJ-sgfp* with pMGPII; $Cam^r$ | This work |
| ColPBP1iRodA-sGFP | COL Δ*pbpA spa::*$P_{spac}$*pbpA rodA::rodA-sgfp* with pMGPII; $Cam^r$ | This work |
| ColΔ*rodA*sGFP-PBP3 | COL Δ*rodA pbpC::sgfp-pbpC* | This work |
| ColΔ*rodA*pRodA-sGFP | COL Δ*rodA* with pCNX-rodAsgfp; $Kan^r$ | This work |
| ColΔ*rodA*pRodA$^{W111A}$-sGFP | COL Δ*rodA* with pCNX-rodA$^{W111A}$sgfp; $Kan^r$ | This work |

| Strains and plasmids | Description | Source or reference |
|---|---|---|
| ColΔ*rodA*pRodA$^{D286A}$-sGFP | COL Δ*rodA* with pCNX-rodA$^{D286A}$sgfp; Kan$^{r}$ | This work |
| ColΔ*pbpC*RodA-sGFP | COL Δ*pbpC rodA*::*rodA-sgfp* | This work |
| ColΔ*pbpC*RodA-sGFPpPBP3 | COL Δ*pbpC rodA*::*rodA-sgfp* with pCNX-pbpC; Kan$^{r}$ | This work |
| **Plasmids** | | |
| pMGPII | *S. aureus* replicative plasmid containing *lacI* gene; Amp$^{r,}$ Cam$^{r}$ | 5 |
| pTRC99a-P7 | Plasmid containing *sgfp-p7* gene, Amp$^{r}$ | 6 |
| pROD17 | Plasmid containing *mCherry* gene | D. Sherratt |
| pMAD | *E. coli-S. aureus* shuttle vector with a thermosensitive origin of replication for Gram positive bacteria; Amp$^{r,}$ Ery$^{r}$, *lacZ* | 7 |
| pSKP1S314A | pSK5632 vector with *pbpA*$^{S314A}$ copy, Amp$^{r}$, Cam$^{r}$ | 8 |
| pBCBAJ001 | pMAD containing the 3' end and the downstream region of ezrA; Amp$^{r}$, Erm$^{r}$ | 9 |
| pMAD-Δ*ftsW* | pMAD derivative containing up-and downstream regions of *ftsW;* Amp$^{r,}$ Ery$^{r}$ | This work |
| pMAD-Δ*pbpA* | pMAD derivative containing upstream and 3' end regions of *pbpA;* Amp$^{r,}$ Ery$^{r}$ | This work |
| pMAD-Δ*pbpC* | pMAD derivative containing up-and downstream regions of *pbpC;* Amp$^{r,}$ Ery$^{r}$ | This work |

| Strains and plasmids | Description | Source or reference |
| --- | --- | --- |
| pΔpbpD | pMAD derivative containing up-and downstream regions of *pbpD;* Amp$^{r,}$ Ery$^{r}$ | 10 |
| pMAD-Δ*rodA* | pMAD derivative containing up-and downstream regions of *rodA;* Amp$^{r,}$ Ery$^{r}$ | This work |
| pMAD-pbpATP | pMAD derivative containing Pspac and full-length *pbpA*$^{S314A}$*;* Amp$^{r,}$ Ery$^{r}$ | This work |
| pMAD-ezrAsgfp | pMAD derivative containing an *ezrA-sgfp* fusion and the downstream region of *ezrA;* Amp$^{r,}$ Ery$^{r}$ | This work |
| pMAD-ftsWsgfp | pMAD derivative containing an *ftsW-sgfp* fusion and the downstream region of *ftsW;* Amp$^{r,}$ Ery$^{r}$ | 4 |
| pMAD-murJsgfp | pMAD derivative containing a *murJ-sgfp* fusion and the downstream region of *murJ;* Amp$^{r}$ Ery$^{r}$ | 4 |
| pMAD-sgfpPbp1 | pMAD derivative containing an *sgfp-pbpA* fusion and the upstream region of *pbpA;* Amp$^{r,}$ Ery$^{r}$ | 4 |
| pMAD-sgfpPbp3 | pMAD derivative containing a *sgfp-pbpC* fusion and the upstream region of *pbpC;* Amp$^{r,}$ Ery$^{r}$ | 4 |
| pMAD-rodAsgfp | pMAD derivative containing a *rodA-sgfp* fusion and the downstream region of *rodA;* Amp$^{r}$ Ery$^{r}$ | 4 |
| pMAD-rodAmCh | pMAD derivative containing a *rodA-mCherry* fusion and the downstream region of *rodA;* Amp$^{r}$ Ery$^{r}$ | This work |
| pBCB13 | pMAD derivative with up- and downstream regions of the *spa* locus and P*spac-lacI*, Amp$^{r}$, Ery$^{r,}$ *lacZ* | 11 |
| pBCB13-ftsW | pBCB13 derivative containing *ftsW* gene under the control of IPTG-inducible P*spac* promoter *;* Amp$^{r}$, Ery$^{r}$ | This work |
| pBCB13-pbpA | pBCB13 derivative containing *pbpA* gene under the control of IPTG-inducible P*spac* promoter*;* Amp$^{r}$, Ery$^{r}$ | This work |
| pBCB13-sgfppbpA | pBCB13 derivative containing an *sgfp-pbpA* fusion under the control of IPTG-inducible P*spac* promoter*;* Amp$^{r}$, Ery$^{r}$ | This work |

| Strains and plasmids | Description | Source or reference |
|---|---|---|
| pCNX | Shuttle vector containing a cadmium inducible P*cad* promoter; $Amp^r$, $Kan^r$ | 12 |
| pCNX-pbpC | pCNX derivative containing *pbpC* gene; $Amp^r$, $Kan^r$ | This work |
| pCNX-ftsWsgfp | pCNX derivative containing an *ftsW-sgfp* fusion; $Amp^r$, $Kan^r$ | This work |
| pCNX-ftsW$^{W121A}$sgfp | pCNX derivative containing an *ftsW*$^{W121A}$*-sgfp* fusion ; $Amp^r$, $Kan^r$ | This work |
| pCNX-ftsW$^{D287A}$sgfp | pCNX derivative containing an *ftsW*$^{D287A}$*-sgfp* fusion; $Amp^r$, $Kan^r$ | This work |
| pCNX-ftsWmCh | pCNX derivative containing an *ftsW-mCherry* fusion; $Amp^r$, $Kan^r$ | This work |
| pCNX-murJmCh | pCNX derivative containing an *murJ-mCherry* fusion; $Amp^r$, $Kan^r$ | This work |
| pCN-ftsZ$^{55\text{-}56}$sGFP | pCNX derivative containing an *ftsZ-sgfp* fusion; $Amp^r$, $Kan^r$ | 4 |
| pCNX-rodAsgfp | pCNX derivative containing a *rodA-sgfp* fusion; $Amp^r$, $Kan^r$ | This work |
| pCNX-rodA$^{W111A}$sgfp | pCNX derivative containing a *rodA*$^{W111A}$*-sgfp* fusion; $Amp^r$, $Kan^r$ | This work |
| pCNX-rodA$^{D286A}$sgfp | pCNX derivative containing a *rodA*$^{D286A}$*-sgfp* fusion; $Amp^r$, $Kan^r$ | This work |
| pCNX-rodAmCh | pCNX derivative containing a *rodA-mCherry* fusion; $Amp^r$, $Kan^r$ | This work |
| pCNX-sgfppbpA | pCNX derivative containing an *sgfp-pbpA* fusion; $Amp^r$, $Kan^r$ | This work |
| pCNX-sgfppbpC | pCNX derivative containing an *sgfp-pbpC* fusion; $Amp^r$, $Kan^r$ | This work |
| pCNX-mChsgfp | pCNX derivative containing an *mCherry-sgfp* fusion; $Amp^r$, $Kan^r$ | This work |

| Strains and plasmids | Description | Source or reference |
|---|---|---|
| pCNX-mChsgfpTM | pCNX derivative containing an *mCherry-sgfp-TM* fusion; $Amp^r$, $Kan^r$ | This work |
| pCNX-WP1 | pCNX derivative containing an *ftsW-mCherry* and *sgfp-pbpA* fusion; $Amp^r$, $Kan^r$ | This work |
| pCNX-JP1 | pCNX derivative containing a *murJ-mCherry* and *sgfp-pbpA* fusion; $Amp^r$, $Kan^r$ | This work |
| pCNX-AP1 | pCNX derivative containing a *rodA-mCherry* and *sgfp-pbpA* fusion; $Amp^r$, $Kan^r$ | This work |
| pCNX-AP3 | pCNX derivative containing a *rodA-mCherry* and *sgfp-pbpC* fusion; $Amp^r$, $Kan^r$ | This work |

### Supplementary Table 2 – Oligonucleotides used in this study

| Primer No. | Primer Name | Sequence (5'-3') |
|---|---|---|
| **202** | pBCB13PBP1Rev | GCAGCTCGAGTTAGTCCGACTTATCCTTGTCAGTTTTAC |
| **1285** | Pspac_pDH88_P1_EcoRI | GCTGAATTCTTCTACACAGCCCAGTCCAGAC |
| **1294** | Up spa_P1_BamHI | TGAGGATCCCTGGTTCAGTTGTAAATAACAATAC |
| **1297** | DOWN spa_P4_NcoI | TGCAGTCCATGGGTTAGAGCTCTCAATAATTTAAAAAAGC |
| **1387** | PBP1fppBCB13FWP3 | GGTGGAGGAGGTTCTGGTGGAGGAGGTTCTATGGCGAAGCAAAAAATTAAAAT |
| **2949** | sfGFP_Cterm_P1_NheI | GCATGGCTAGCATGAGTAAAGGAGAAGAACTTTTC |
| **3112** | XmaI-RBS-GFP(P7) for | GCGCCCGGGAAAAAATAAGGAGGAAAAAAAATGAGTAAAGGAGAAGAACTTTTC |
| **3211** | GFP(P7)-10aa linker revx | AGAACCTCCTCCACCAGAACCTCCTCCACCGTCGACTTTGTATAGTTCATCCATG |
| **3350** | GFP(P7)-TAA-XhoI rev | CCGGCTCGAGTTATTTGTATAGTTCATCCATG |
| **3490** | PBP1-BamHI rev | GCCGGATCCTTAGTCCGACTTATCCTTGTC |
| **3599** | BamHI-RBS for | GCCGGATCCAAAAAATAAGGAGGAAAAAAAATG |
| **3600** | PBP1-EcoRI rev | GCCGAATTCTTAGTCCGACTTATCCTTGTC |
| **3806** | BglII-SACOL1192(last183bp) for | CCGAGATCTCAGGTGTTCCAAGAATATG |
| **3809** | GFP(P7)-10aa linker rev | AGAGCCACCTCCGCCAGAACCGCCTCCACCGTCGACTTTGTATAGTTCATCC |
| **4041** | C-GFP-KpnI Rev | CCGGTACCTTATTTGTATAGTTCATCCATGCCATGTG |
| **4235** | Pspac-RBS Rev | CATTTTTTTTCCTCCTTATTTTTTCCCGGGAAAAGCTTAATTGTTATCC |
| **4236** | RBS-pbpA For | GAAAAAATAAGGAGGAAAAAAAATGGCGAAGCAAAAAATTAAAATT |
| **4470** | SmaI-RBS-pbpA For | GCGCCCGGGAAAAAATAAGGAGGAAAAAAAATGGCGAAGCAAAAAATTAAAATT |
| **5187** | TMpbp2_Xma_Rev | TCGGTACCCGGGTTAAGCAAACAATAAGATACCTAG |
| **5440** | BamHI-RBS-ATG For | CGCGGATCCAAAAAATAAGGAGGAAAAAAAATG |
| **5668** | PBP3-EcoRI Rev | GCCGAATTCTTATTTGTCTTTGTCTTTATTTTTATC |
| **5671** | CFP-EcoRI Rev | GCCGAATTCTTACTTGTACAGCTCGTCCATGCC |
| **5706** | BamHI-RBS-1122 For | CGCGGATCCAAAAAATAAGGAGGAAAAAAAATGAAGAATTTTAGAAGTATTTTAC |
| **5707** | BamHI-RBS-2075 For | CGCGGATCCAAAAAATAAGGAGGAAAAAAAATGAATTATTCATCTCGTCAACAG |

| | | |
|---|---|---|
| **5708** | BamHI-RBS-1804 For | CGCGGATCCAAAAAATAAGGAGGAAAAAAAATGAGTGAAAGTAAAGAAATGGTG |
| **5772** | 1193-pbpA1.9kb Rev | GATTGTGCTTTTATTTAATTTTTTGCTTCGCCATTAC |
| **5773** | 1193-pbpA1.9kb For | CAAAAAATTAAATAAAAGCACAATCTATAAAAGCAG |
| **5774** | 250dwnpbpA-BamHI Rev | CGCGGATCCCCACTAATATATTATCAATTTTTC |
| **5941** | CFP-KpnI Rev | GCCGGTACCTTACTTGTACAGCTCGTCCATGCC |
| **6052** | CFP-NotI Rev | GGGGCGGCCGCTTACTTGTACAGCTCGTCCATGCC |
| **6053** | NotI-RBS For | GGGGCGGCCGCAAAAAATAAGGAGGAAAAAAAATG |
| **6398** | mCh-KRSGSGGS Rev | GGAGCCACCAGAACCAGATCTTTTCTTGTACAGCTCGTCCATGCCACC |
| **6399** | KRSGSGGS-GFP For | GAAAAGATCTGGTTCTGGTGGCTCCAGTAAAGGAGAAGAACTTTTCAC |
| **6404** | (RBS)-ATG-GFP For | GGAGGAAAAAAAATGAGTAAAGGAGAAGAACTTTTC |
| **6400** | GGGSGGGGS-TM2 For | GGAGGCGGTTCTGGCGGAGGTGGCTCTACGGAAAACAAAGGATCTTCTCAG |
| **6407** | (RBS)-ATG-mCh For | GGAGGAAAAAAAATGGCTATCATTAAAGAGTTCATG |
| **4179** | P1 SACO1122i_spa | ACTCCCGGGATCATTTGAAGTATAAATTG |
| **4180** | P2 SACO1122i_spa | AGTCTCGAGATCTCGTACTAAATATTGG |
| **3049** | P1 SACOL1122 KO | AGTCCCGGGGTCGTACTATCTAATGTTATAGG |
| **3050** | P2 SACOL1122 KO | TTGGCTAGTATTTTTTTAATATTCAGTCATCCAATTC |
| **3051** | P3 SACOL1122 KO | ATTGTAGAATTGGATGACTGAATATTAAAAAAATACTAGCC |
| **3052** | P4 SACOL1122 KO | ACTGGATCCTGCTACATCAATGATACGCTC |
| **3056** | P1 SACOL2075 KO | ATTCCCGGGCATAGAACTTGCAGCTGACAATACACC |
| **3057** | P2 SACOL2075 KO | TATTTGAAACTCAAAATAGTTTAAATTATGCAAACTCCTTTTATACTCAC |
| **3058** | P3 SACOL2075 KO | AGTATAAAAGGAGTTTGCATAATTTAAACTATTTTGAGTTTC |
| **3059** | P4 SACOL2075 KO | ATAGGATCCGTAAGTCAATTGACAGAACGC |
| **2284** | P1_SACOL2075_pMaD | AATCCCGGGTATCAGACAAATTTTTTATTACATTTTAGGTGC |
| **5665** | 2075-mCh_P4 | AGAGCCACCTCCGCCAGAACCGCCTCCACCATTACTTTTT GGATGGTATAAATC |
| **5546** | 2075-mCh_P3 | GTACAAGTAATTTAAACTATTTTGAGTTTC |
| **2287** | P6_SACOL2075_pMAD | ATTGGATCCATGAAGGAGTGAATGCTATGACT |
| **5599** | 10aa-mCh For | GGCGGTTCTGGCGGAGGTGGCTCTGCTATCATTAAAGAGTTCATGCGC |
| **5545** | 2075-mCh_P2 | ATAGTTTAAATTACTTGTACAGCTCGTCCATG |
| **4041** | C-GFP-KpnI Rev | CCGGTACCTTATTTGTATAGTTCATCCATGCCATGTG |

| | | |
|---|---|---|
| **5603** | GFPSTOPP2_Eco | CGCCGAATTCTTATTTGTATAGTTCATCCATGCC |
| **5609** | SACOL2075_P1_BamHI | CGCGGATCCTGAGTATAAAAGGAGTTTGC |
| **5547** | 1122_W121A_P1 | ATATTAATGGTTCTAAAAGTGCAATAAACTTAGGATTTATGAAC |
| **5548** | 1122_W121A_P2 | TTCATAAATCCTAAGTTTATTGCACTTTTAGAACCATTAATATC |
| **5553** | 1122_D287A_P1 | ATTTACCAGAACCACATACAGCATTTATTTTTGCAATTATTTG |
| **5554** | 1122_D287A_P2 | CAAATAATTGCAAAAATAAATGCTGTATGTGGTTCTGGTAAATAG |
| **5559** | 2075_W111A_P1 | TTATCAATGGTGCCAAAAGTGCATACACGTTTGGCCCTATCAG |
| **5560** | 2075_W111A_P2 | CTGATAGGGCCAAACGTGTATGCACTTTTGGCACCATTGATAATAG |
| **5565** | 2075_D286A_P1 | ATATACCTGAAAATCATACTGCATTTATCTTTTCAGTGATTTG |
| **5566** | 2075_D286A_P2 | CCAATCACTGAAAAGATAAATGCAGTATGATTTTCAGGTATATAAAC |
| **2587** | pPBP3-KO-P1 | ATCGAGAATTCATGCACATTTGGTCAG |
| **2588** | pPBP3-KO-P2 | TCGTCAGGTTAAATTAACTACCTACCTC |
| **2589** | pPBP3-KO-P3 | AGAGGTAGGTAGTTAATTTAACCTGACG |
| **2590** | pPBP3-KO-P4 | AGTCGGATCCCTATGCTGCTTGAT |
| **5704** | BamHI-RBS-PBP3 For | CGCGGATCCAAAAAATAAGGAGGAAAAAAATTGTTAAAAAGACTAAAAGAAAAAT |

Underlined sequences correspond to restriction sites

Bibliography


1 Monk, I. R., Shah, I. M., Xu, M., Tan, M. W. & Foster, T. J. Transforming the untransformable: application of direct transformation to manipulate genetically *Staphylococcus aureus* and *Staphylococcus epidermidis*. *mBio* **3**, doi:10.1128/mBio.00277-11 (2012).

2 Nair, D. *et al.* Whole-genome sequencing of *Staphylococcus aureus* strain RN4220, a key laboratory strain used in virulence research, identifies mutations that affect not only virulence factors but also the fitness of the strain. *J Bacteriol* **193**, 2332-2335, doi:10.1128/jb.00027-11 (2011).

3 Gill, S. R. *et al.* Insights on evolution of virulence and resistance from the complete genome analysis of an early methicillin-resistant *Staphylococcus aureus* strain and a biofilm-producing methicillin-resistant *Staphylococcus epidermidis* strain. *J. Bacteriol.* **187**, 2426-2438, doi:10.1128/JB.187.7.2426-2438.2005 (2005).

4 Monteiro, J. M. *et al.* Peptidoglycan synthesis drives an FtsZ-treadmilling-independent step of cytokinesis. *Nature* **554**, 528-532, doi:10.1038/nature25506 (2018).

5 Pinho, M. G., Filipe, S. R., de Lencastre, H. & al., e. Complementation of the essential peptidoglycan transpeptidase function of penicillin-binding protein 2 (PBP2) by the drug resistance protein PBP2A in *Staphylococcus aureus*. *J. Bacteriol.* **183**, 6525-6531, doi:10.1128/JB.183.22.6525-6531.2001 (2001).

6 Fisher, A. C. & DeLisa, M. P. Laboratory evolution of fast-folding green fluorescent protein using secretory pathway quality control. *PLoS ONE* **3**, e2351, doi:10.1371/journal.pone.0002351 (2008).

7 Arnaud, M., Chastanet, A. & Debarbouille, M. New vector for efficient allelic replacement in naturally nontransformable, low-GC-content, gram-positive bacteria. *Appl. Environ. Microbiol.* **70**, 6887-6891, doi:10.1128/AEM.70.11.6887-6891.2004 (2004).

8 Pereira, S. F. F., Henriques, A. O., Pinho, M. G., de Lencastre, H. & Tomasz, A. Evidence for a dual role of PBP1 in the cell division and cell separation of *Staphylococcus aureus*. *Molecular Microbiology* **72**, 895-904, doi:10.1111/j.1365-2958.2009.06687.x (2009).

9 Jorge, A. M., Hoiczyk, E., Gomes, J. P. & Pinho, M. G. EzrA Contributes to the Regulation of Cell Size in *Staphylococcus aureus*. *PLoS ONE* **6**, e27542, doi:10.1371/journal.pone.0027542 (2011).

10 Atilano, M. *et al.* Teichoic acids are temporal and spatial regulators of peptidoglycan cross-linking in *Staphylococcus aureus*. *Proc. Natl. Acad. Sci. U S A* **107**, 18991-18996, doi:10.1073/pnas.1004304107 (2010).

11 Pereira, P., Veiga, H., Jorge, A. & Pinho, M. Fluorescent Reporters for Studies of Cellular Localization of Proteins in *Staphylococcus aureus*. *Appl. Environ. Microbiol.* **76**, 4346-4353, doi:10.1128/AEM.00359-10 (2010).

12 Monteiro, J. M. *et al.* Cell shape dynamics during the staphylococcal cell cycle. *Nat Commun* **6**, 8055, doi:10.1038/ncomms9055 (2015).
