## Supplementary methods for "SEDS-bPBP pairs direct Lateral and Septal Peptidoglycan Synthesis in *Staphylococcus aureus*"

### MATERIALS AND METHODS

#### Bacterial growth conditions

Strains and plasmids used in this study are listed in Supplementary Table 1. *E. coli* strains were grown in Luria–Bertani broth (Difco) or Luria–Bertani agar (Difco) at 37°C. *S. aureus* strains were grown in tryptic soy broth (TSB, Difco) or on tryptic soy agar (TSA, Difco) at 37°C. When necessary, culture media was supplemented with antibiotics (ampicillin 100 µg $ml^{-1}$, kanamycin 50 µg $ml^{-1}$, neomycin 50 µg $ml^{-1}$, erythromycin 10 µg $ml^{-1}$ or chloramphenicol 10 µg $ml^{-1}$), with 100 µg $ml^{-1}$ 5-Bromo-4-chloro-3-indolyl β-D-galactopyranoside (X-gal, from Apollo Scientific), with 0.5 mM (ColFtsWi) or 0.01 mM (ColPBP1i) Isopropyl β-D-1-thiogalactopyranoside (IPTG, Apollo Scientific) or with 0.1 µM of cadmium chloride (Sigma-Aldrich).

#### Construction of *S. aureus* strains

Primers used in this study are listed in Supplementary Table 2. Plasmids were initially constructed in *E. coli,* then introduced in *S. aureus* RN4220 strain by electroporation, as previously described[1] and finally transduced to the strain of interest using phage 80α[2]. For the construction of ColFtsWi and ColPBP1i, a second copy of each gene was first introduced at the ectopic *spa* locus, under the control of an IPTG-inducible P*spac* promoter, prior to the deletion of the native loci copies. Briefly, a DNA fragment containing an RBS and the *ftsW* or *pbpA* gene was amplified by PCR from *S. aureus* strain COL with primers 4179/4180, and 4470/202, respectively. Each PCR fragment was digested with SmaI/XhoI and cloned into pBCB13, downstream of the P*spac* promoter, giving pBCB13-ftsW and pBCB13-pbpA. Integration/excision was performed in COL as previously described[3], resulting in ColFtsWhigh and ColPBP1high.

In order to delete the native copies of *ftsW*, *rodA*, and *pbpC* (encoding PBP3), upstream and downstream regions of each gene of interest were amplified from COL using primers 3049/3050 and 3051/3052 (for *ftsW)*, 3056/3057 and 3058/3059 (for *rodA*), and 2587/2588 and 2589/2590 (for *pbpC*). In each case, the two fragments were joined by overlap PCR using primers 3049/3052, 3056/3059 and 2587/2590, respectively, digested with SmaI/BamHI (*ftsW* and *rodA*) or EcoRI/BamHI (*pbpC*) and cloned into pMAD, originating pMAD-Δ*ftsW*, pMAD-Δ*rodA* and pMAD-Δ*pbpC*. Integration/excision of pMAD-Δ*rodA* and pMAD-Δ*pbpC* in COL resulted in strains ColΔ*rodA* and ColΔ*pbpC*. Plasmid pMAD-Δ*rodA* was also introduced into ColsGFP-PBP3[4] and ColΔ*pbpC*, and integration/excision generated ColΔ*rodA*sGFP-PBP3 and

ColΔ*rodA*Δ*pbpC*, respectively. Integration/excision of pMAD-rodAsgfp in ColΔ*pbpC* resulted in ColΔ*pbpC*RodA-sGFP.

Despite several attempts, deletion of the full *pbpA* gene (encoding PBP1) was unsuccessful. In case we were affecting the expression of the essential *mraY* gene (290bp downstream of *pbpA*), we instead constructed pMAD-Δ*pbpA* to include a stop codon followed by the last 340bp of *pbpA* using primers 3806/5772 and 5773/5774. Overlap PCR was performed with primers 3806/5774 and the full fragment was digested with BglII/BamHI and cloned into pMAD.

Plasmids pMAD-Δ*ftsW* and pMAD-Δ*pbpA* were introduced into ColFtsWhigh and ColPBP1high, and integration/excision in the presence of IPTG resulted in deletion of the native genes, originating ColFtsWlow and ColPBP1low. Plasmid pMGPII, encoding the P*spac* repressor LacI was transduced into these strains and into COL, giving rise to ColFtsWi, ColPBP1i and ColpMGPII.

For the construction of ColPBP1TP, a *pbpA* allele encoding the TP point mutation S314A (by substitution of *tca* for *gca* at codon 314) was placed under the control of the P*spac* promoter by first amplifying the P*spac* promoter and an RBS from pBCB13 using primers 1285/4235, and the *pbpA* gene encoding for $PBP1^{S314A}$ using primers 4236/3490 from pSKP1S314A. The two fragments were joined by overlap PCR using primers 1285/3490, digested with EcoRI/BamHI and cloned into pMAD, to give pMAD-pbpATP. Integration/excision was performed in COL in the presence of 0.5 mM IPTG.

For localisation studies of EzrA, sGFP was amplified from pTRC99a-P7 using primers 2949/3350, digested with NheI/XhoI and cloned into pBCBAJ001, giving pMAD-ezrAsgfp, which was transduced to ColFtsWi and ColPBP1i. Integration/excision resulted in the introduction of a 3´ *sgfp* fusion to the native copy of *ezrA*, giving strains ColFtsWiEzrA-sGFP and ColPBP1iEzrA-sGFP, respectively.

For construction of *pbpD* deletion strains, plasmid pΔpbpD was transduced into ColFtsWi, ColPBP1i, ColFtsWiEzrA-sGFP and ColPBP1iEzrA-sGFP, and following integration/excision generated strains ColFtsWiΔ*pbpD*, ColPBP1iΔ*pbpD*, ColFtsWiΔ*pbpD*EzrA-sGFP and ColPBP1iΔ*pbpD*EzrA-sGFP, respectively.

For localisation of FtsZ, pCN-$ftsZ^{55-56}$sGFP was transduced into ColFtsWi and ColPBP1i, generating ColFtsWipFtsZ-sGFP and ColPBP1ipFtsZ-sGFP, respectively. For localisation studies in ColPBP1i, transduction and integration/excision of pMAD-ftsWsgfp and pMAD-murJsgfp plasmids, gave rise to ColPBP1iFtsW-sGFP and ColPBP1iMurJ-sGFP, respectively.

For colocalisation studies with FtsW-mCherry, plasmids pMAD-sgfpPbp1 and pMAD-sgfpPbp3 were transduced into ColFtsW-mCherry and following integration/excision gave strain ColWP1 and ColWP3, respectively. For colocalisation studies with RodA-mCherry, this fusion was first introduced into COL background. Briefly, the final 1051 bp of the *rodA* gene, excluding the stop codon, and its downstream region were amplified using primers 2284/5665 and 5546/2287, respectively. The *mCherry* gene was amplified from pROD17 plasmid using primers 5599/5545. Fragments were joined by overlap PCR using primers 2284/2287, digested with SmaI/BamHI and cloned into pMAD vector, originating pMAD-rodAmCh. Integration and excision in COL gave rise to ColRodA-mCherry. The *rodA-mCherry* fusion was then amplified from ColRodA-mCherry using primers 5707/5941, digested with BamHI/KpnI and cloned into pCNX, resulting in pCNX-rodAmCh. This plasmid was transduced into COL, ColsGFP-PBP1 and ColsGFP-PBP3, giving ColpRodA-mCherry, ColP1pA and ColP3pA, respectively.

For inducible expression of fluorescent fusions, *ftsW-mCherry* and *murJ-mCherry* were amplified from strains ColFtsW-mCherry and ColMurJ-mCherry using primers 5706/5671 and 5708/5671, digested with BamHI/EcoRI and cloned into pCNX, resulting in pCNX-ftsWmCh and pCNX-murJmCh, respectively. These plasmids and empty pCNX vector were introduced into COL, giving ColpFtsW-mCherry, ColpMurJ-mCherry and ColpCNX, respectively.

For inducible expression of sGFP-PBP1, *sgfp-pbpA* was first cloned into plasmid pBCB13, before amplification and introduction into pCNX. For construction of pBCB13-sgfppbpA, a fragment containing *sgfp* was amplified from pTRC99a-P7 using primers 3112/3211 and the *pbpA* gene was amplified from COL using primers 1387/202. The two fragments were joined by overlap PCR using primers 3112/202, digested with SmaI/XhoI and cloned into pBCB13. Primers 3599/3600 were then used to amplify the *sgfp-pbpA* fragment, digested with BamHI/EcoRI and cloned into pCNX, resulting in pCNX-sgfppbpA. This plasmid was transduced into COL, giving ColpsGFP-PBP1.

For inducible expression of sGFP-PBP3, primers 6404/5668 were used to amplify *sgfp-pbpC* from ColsGFP-PBP3, introducing an RBS at the 5' end. This PCR was re-amplified using primers 5440/5668 in order to add a BamHI restriction site, and following digestion with BamHI/EcoRI, was cloned into pCNX, resulting in pCNX-sgfppbpC. Transduction of this plasmid into COL gave ColpsGFP-PBP3.

For inducible expression of PBP3, primers 5704/5668 were used to amplify *pbpC* from COL, digested with BamHI/EcoRI, and cloned into pCNX, resulting in pCNX-pbpC. Transduction of this plasmid into ColΔ*pbpC*RodA-sGFP, generated ColΔ*pbpC*RodA-sGFPpPBP3.

For complementation experiments, sGFP fusions to wild-type and point mutants of FtsW and RodA were expressed from pCNX. Briefly, *ftsW-sgfp* and *rodA-sgfp* fragments were amplified from ColFtsW-sGFP and ColRodA-sGFP using primers 4179/5603 and 5609/4041, digested with SmaI/EcoRI or BamHI/KpnI and cloned into pCNX vector, originating pCNX-ftsWsgfp and pCNX-rodAsgfp. In FtsW, amino acid exchange was performed by altering *ftsW* codon 121 from *tgg* to *gca* (W121A) and codon 287 from *gat* to *gca* (D287A). Alterations were performed by amplifying pCNX-ftsWsgfp with primers 5547/5548 or 5553/5554, originating plasmids pCNX-ftsW$^{W121A}$sgfp and pCNX-ftsW$^{D287A}$sgfp. These plasmids, pCNX and pCNX-ftsWsgfp were introduced into ColFtsWi, resulting in ColFtsWipFtsW$^{W121A}$-sGFP, ColFtsWipFtsW$^{D287A}$-sGFP, ColFtsWipCNX and ColFtsWipFtsW-sGFP, respectively.

For RodA, amino acid exchange was performed by altering *rodA* codon 111 from *tgg* to *gca* (W111A) and codon 286 from *gac* to *gca* (D286A). Alterations were performed by amplifying pCNX-rodAsgfp with primers 5559/5560 or 5565/5566, originating plasmids pCNX-rodA$^{W111A}$sgfp and pCNX-rodA$^{D286A}$sgfp. These plasmids, and pCNX-rodAsgfp, were introduced into ColΔ*rodA* resulting in ColΔ*rodA*pRodA$^{W111A}$-sGFP, ColΔ*rodA*pRodA$^{D286A}$-sGFP and ColΔ*rodA*pRodA-sGFP, respectively.

For FLIM and seFRET experiments, plasmids coexpressing fluorescent fusions were designed. In general, mCherry fluorescent protein fusions were amplified with an RBS and flanked with BamHI and NotI restriction sites, and sGFP fluorescent protein fusions were amplified with an RBS and flanked with NotI and EcoRI sites. Digested PCR products were ligated to BamHI/EcoRI-digested pCNX, resulting in an RBS and mCherry fusion, followed by a second RBS and sGFP fusion placed under a cadmium inducible promoter. Specifically, *ftsW-mCherry*, *murJ-mCherry* and *rodA-mCherry* were amplified using primers 5706/6052, 5708/6052 and 5707/6052 from ColFtsW-mCherry, ColMurJ-mCherry and ColRodA-mCherry, respectively. GFP fusions *sgfp-pbpA* and *sgfp-pbpC* were amplified using primers 6053/3600 and 6053/5668 from ColsGFP-PBP1 and ColsGFP-PBP3, respectively. Digestion and ligation gave rise to plasmids pCNX-WP1, pCNX-JP1, pCNX-AP1 and pCNX-AP3, and transduction into COL resulted in strains ColpWP1, ColpJP1, ColpAP1 and ColpAP3, respectively.

As a positive membrane control for FLIM, a fusion between *mCherry*, *sgfp* and a fragment of *pbpB* encoding its transmembrane domain (amino acids 2 to 34) was cloned into pCNX. Primers 6407/6398 were used to amplify *mCherry* from ColFtsW-mCherry, *gfp* was amplified from pTRC99a-P7 using primers 6399/3809, and the *pbpB* fragment was amplified from COL using primers 6400/5187. These three fragments were joined by overlap PCR using primers 3599/5187, digested with BamHI/SmaI and cloned into pCNX, resulting in pCNX-mChsgfpTM. Transduction into COL gave ColpmCherry-sGFP-TM.

As a positive cytoplasmic control for seFRET, a fusion between *mCherry* and *sgfp* was cloned into pCNX. Primers 6407/6398 were used to amplify *mCherry* from ColFtsW-mCherry and *gfp* was amplified from pTRC99a-P7 using primers 6399/4041. These two fragments were joined by overlap PCR using primers 3599/4041, digested with BamHI/KpnI and cloned into pCNX, resulting in pCNX-mChsgfp. Transduction into COL gave ColpmCherry-sGFP.

**Next Generation Sequencing (NGS)**

Chromosomal DNA was purified from ColPBP1TP using a standard phenol-chloroform extraction technique[5]. DNA was sequenced at Instituto Gulbenkian de Ciência, Oeiras, Portugal, using the Illumina MiSeq system, producing 300 bp paired end reads with >100 x average coverage. Sequence assembly was performed with SeqMan NGen 12 software (DNASTAR, Inc) using the COL genome, and with the exception of the PBP1 TP point mutation introduced, no single nucleotide polymorphisms (SNPs) that could be confirmed by Sanger sequencing were identified.

**Growth curves**

Overnight cultures were back-diluted to $OD_{600nm}$ 0.02 in TSB and grown for 8 hours with $OD_{600nm}$ measurements taken every hour. In the case of ColFtsWi, ColPBP1i and their derivatives, overnight cultures were washed four times with TSB and back-diluted to $OD_{600}$ of 0.02 in TSB with or without IPTG. At $OD_{600nm}$ of approximately 1, cells were similarly washed and back-diluted to $OD_{60onm}$ of 0.05. $OD_{600nm}$ measurements were taken every hour.

**Minimum Inhibitory Concentration (MIC) assays**

Moenomycin MIC was determined using broth microdilution in sterile 96-well plates, where overnight cultures were diluted to a cell density of ~$5x10^3$ CFU $ml^{-1}$ in wells containing two-fold dilutions of moenomycin. Plates were incubated aerobically for 24hr at 37°C, and endpoint growth was assessed visually. All assays were performed in triplicate.

**HPLC analysis of muropeptides**

Peptidoglycan purification from mid-exponential phase cultures was performed as previously described[6]. Preparation of muropeptides was performed by digestion with mutanolysin (Sigma) and reduction with sodium borohydride (Sigma), before analysis by reverse-phase HPLC using a Hypersil ODS column (Thermo Electron Corporation).

***S. aureus* imaging by fluorescence microscopy**

Structured Illumination Microscopy (SIM) imaging was performed using an Elyra PS.1 microscope (Zeiss) with a Plan-Apochromat 63x/1.4 oil DIC M27 objective. SIM images were acquired using three or five grid rotations, with 34 µm grating period for the 561 nm laser

(100 mW), 28 µm period for 488 nm laser (100 mW) and 23 µm period for 405 nm laser (50 mW), and captured using a Pco.edge 5.5 camera. Images were reconstructed using ZEN software (black edition, 2012, version 8.1.0.484) based on a structured illumination algorithm, using synthetic, channel specific optical transfer functions and noise filter settings ranging from −6 to −8.

Epifluorescence microscopy for colocalisation studies and fluorescence ratio analysis was performed using a Zeiss Axio Observer microscope with a Plan-Apochromat 100x/1.4 oil Ph3 objective. Images were acquired with a Retiga R1 CCD camera (QImaging) using Metamorph 7.5 software (Molecular Devices).

For fluorescence microscopy experiments, *S. aureus* cultures were grown to mid-exponential phase ($OD_{600nm}$ of 0.6-1.0) prior to pelleting. Cells were suspended in phosphate buffer saline (PBS) and placed on a thin layer of agarose (1.2% in PBS). For timelapse microscopy, cells were suspended in TSB and spotted on 1.2% agarose in 50% TSB/PBS slide, before imaging every 3 or 5 min. In the case of ColFtsWi, ColPBP1i and their derivatives, overnight cultures were back-diluted to an $OD_{600nm}$ of 0.1 and grown for 3 hours in the presence of 0.5 mM and 0.01 mM IPTG, respectively. Cells were washed three times with TSB and grown in the presence or absence of IPTG. Cells were then harvested and prepared for microscopy as described above.

To observe the localisation of peptidoglycan incorporation, *S. aureus* cells were consecutively incubated with 250 µM of fluorescent D-amino-acids HADA, NADA and TDL[7,8] for 10 min, as previously described[9]. Cells were washed with PBS, mounted on a PBS agarose pad and imaged.

To label *S. aureus* membranes, cells were stained with Nile Red (Invitrogen) at a final concentration of 5 µg $ml^{-1}$ for 5 minutes at 37°C, washed with PBS and then placed on PBS agarose pads for imaging. To label *S. aureus* cell wall, cells were stained with a mixture of equal volumes of vancomycin (Sigma) and a BODIPY FL conjugate of vancomycin (Van-FL, Molecular Probes) at a final concentration of 0.8 µg $ml^{-1}$ for 5 minutes at 37°C, washed with PBS and then placed on PBS agarose pads for imaging.

**Microscopy Analysis**

Cell axes and volumes were calculated as previously described[10]. In brief, an ellipse was first fitted to the border of Nile-Red-stained cells, overlaying the fluorescence signal. Measurement of the longer and shorter axes were then measured and subsequently used to calculate the ratio of long/short axis (L/S). These axes were also used to calculate the volume based on a prolate spheroid.

Fluorescence ratio (FR) of septal to membrane signal was measured using the 25% brightest pixels, using our in-house software, eHooke, as previously described[4].
For cell cycle distribution analysis, Nile-Red-stained cells were classified into one of three cell cycle phases, as previously described[10]. Assays were performed in triplicate.

**Transmission Electron Microscopy (TEM)**

Exponentially growing cells of COL and ColPBP1TP were harvested by centrifugation. ColFtsWi and ColPBP1i were initially grown in the presence of IPTG before washing and back-dilution with and without IPTG until mid-exponential phase. Sample preparation was performed essentially as previously described[11]. Briefly, cell pellets were suspended in a primary fixative (2.5% glutaraldehyde + 1% osmium tetroxide in 0.1 M PIPES buffer at pH 7.2) for 1hr at 4°C, with gentle movement. Cells were then washed five times with MilliQ $H_2O$ to remove the fixative and suspended in 3-4% agarose. Small sections of agarose-embedded cells were incubated overnight at 4°C in 0.5% uranyl acetate. The following day, sample were washed with MilliQ $H_2O$ twice and dehydrated using 10 min steps in ethanol (30-100%), anhydrous ice-cold acetone and anhydrous room temperature acetone. Samples were gradually shifted into 100% Spurr's resin and polymerised for 24hrs at 60°C. Ultrathin sections (90nm) were mounted on 200 mesh Cu grids and stained with Reynold's lead citrate. Excess stain was removed with degassed water and TEM imaging at 120kV was performed on a FEI Tecnai 12 microscope, using the Gatan OneView CMOS camera with Digital Micrograph 3.0 software.

**Fluorescence lifetime imaging microscopy (FLIM)**

Cultures were grown to mid-exponential phase in TSB supplemented with 0.1µM cadmium chloride. Cells were pelleted and suspended in PBS, before being spotted on a PBS agarose pad. FLIM measurements were performed by time correlated single photon counting (TCSPC) using a confocal microscope coupled to a multiphoton Titanium-Sapphire laser (Spectra - Physics Mai Tai BB) as the excitation source. FLIM data was acquired during 120 seconds. The excitation wavelength was set to 840 nm and emission light was selected with a dichroic beam splitter with an excitation SP700 short-pass filter and an emission 525±25 nm band-pass filter inserted in front of the photomultiplier. Images were acquired using a Becker and Hickl SPC 830 module. Fluorescence decays in the septum of each cell were calculated by integrating the FLIM data for all pixels of each septum. Fluorescence lifetimes were obtained by analysing the fluorescence decays through a least square iterative re-convolution of decay functions with the instrument response function (IRF) using the software SPCImage (Becker and Hickl). Monoexponential decays were considered in this analysis. Average FRET efficiencies ($E$) in each cell can be determined from $E = 1 - \frac{\tau_{DA}}{\tau_D}$, where $\tau_{DA}$ and $\tau_D$ are the donor fluorescence lifetime in the presence and absence of acceptor, respectively.

**seFRET Quantification.**

COL cells expressing mCherry-sGFP (cytoplasmic tandem, E=12% as determined by FLIM-FRET), sGFP fusion only (donor-only), mCherry fusion only (acceptor-only), both sGFP and mCherry fusions (double), as well as COL cells containing the empty pCNX vector were grown to mid-exponential phase. In order to obtain microscopy images with the five types of cells, a similar number of cells from each strain was mixed together, pelleted, suspended in PBS and placed on a PBS agarose slide. Imaging was performed on a Zeiss Axio Observer.Z1 microscope equipped with a Photometrics CoolSNAP HQ2 camera (Roper Scientific) and a Colibri Light Emitting Diode (LED) illumination system (Zeiss), using ZEN blue software. Slides were imaged for donor signal (300 msec, $ex_{470nm}/em_{502.5\text{-}537.5nm}$), FRET signal (1500 msec, $ex_{470nm}/em_{603.5\text{-}678.5nm}$) and acceptor signal (3000 msec, $ex_{555nm}/em_{603.5\text{-}678.5nm}$) sequentially. FRET efficiency was calculated as described in Chen *et al*[12], by measuring the autofluorescence in each channel of ColpCNX cells, calculating correction factors for channel bleeding and cross-channel excitation using donor-only and acceptor-only cells, and calculating the ratio between sensitized acceptor emission and quenched donor emission due to FRET (G)[13] using the cytoplasmic tandem and the FRET efficiency value determined by earlier FLIM experiments. The G value[13] was used to quantify the FRET efficiency (E) of sets of proteins, using the donor, acceptor and FRET channel measurements in double strain cells.

To perform seFRET quantification on subcellular regions, in-house developed software was used to identify single cells, as previously described[4]. The background for each cell is locally calculated and removed from each channel signal. Additionally, autofluorescence (median signal value of wild type cells) is also subtracted. To define subcellular regions, the software then separates the membrane of each cell by expanding the cell outline inwards. Following membrane identification, the software finds the septum using an isodata algorithm, as this will be the inner region with the most florescence. After this segmentation, the software asks for user input to define which cells correspond to each strain based on the presence/absence of donor and acceptor signals.  Correction factors are taken from the membrane and septum signals, and the E values are then calculated for each cell of strains expressing a pair of proteins, using septal signal only.

Github repository:  **https://github.com/BacterialCellBiologyLab/PyFRET**

**Statistical analysis.**

Statistical analyses were done using GraphPad Prism 6 (GraphPad Software). Two-tailed Mann-Whitney *U* tests were used to evaluate the differences in cellular volume and shape between cell cycle stages, as well as to compare fluorescence ratios between peripheral and septal wall signal intensity. *P* values ≤ 0.05 were considered as significant for all analysis performed and were indicated with asterisks: *$P \leq 0.05$, **$P \leq 0.01$, ***$P \leq 0.001$ and ****P≤0.0001.

**References:**


1 Veiga, H. & Pinho, M. G. Inactivation of the SauI Type I Restriction-Modification System Is Not Sufficient To Generate *Staphylococcus aureus* Strains Capable of Efficiently Accepting Foreign DNA. *Appl. Environ. Microbiol.* **75**, 3034-3038, doi:10.1128/AEM.01862-08 (2009).

2 Oshida, T. & Tomasz, A. Isolation and characterization of a Tn*551*-autolysis mutant of *Staphylococcus aureus*. *J Bacteriol* **174**, 4952-4959 (1992).

3 Pereira, P., Veiga, H., Jorge, A. & Pinho, M. Fluorescent Reporters for Studies of Cellular Localization of Proteins in *Staphylococcus aureus*. *Appl. Environ. Microbiol.* **76**, 4346-4353, doi:10.1128/AEM.00359-10 (2010).

4 Monteiro, J. M. *et al.* Peptidoglycan synthesis drives an FtsZ-treadmilling-independent step of cytokinesis. *Nature* **554**, 528-532, doi:10.1038/nature25506 (2018).

5 Sambrook, J., Fritsch, E. F. & Maniatis, T. *Molecular cloning: a laboratory manual.* 2 edn, (Cold Spring Harbor Laboratory Press, 1989).

6 Filipe, S. R., Tomasz, A. & Ligoxygakis, P. Requirements of peptidoglycan structure that allow detection by the Drosophila Toll pathway. *EMBO Rep* **6**, 327-333, doi:10.1038/sj.embor.7400371 (2005).

7 Kuru, E. *et al.* In situ Probing of Newly Synthesized Peptidoglycan in Live Bacteria with Fluorescent D-Amino Acids. *Angew. Chem. Int Ed. Engl.* **51**, 12519-12523, doi:10.1002/anie.201206749 (2012).

8 Kuru, E., Tekkam, S., Hall, E., Brun, Y. V. & Van Nieuwenhze, M. S. Synthesis of fluorescent D-amino acids and their use for probing peptidoglycan synthesis and bacterial growth in situ. *Nat. Protoc.* **10**, 33-52, doi:10.1038/nprot.2014.197 (2015).

9 Pereira, A. R. *et al.* FtsZ-Dependent Elongation of a Coccoid Bacterium. *mBio* **7**, doi:10.1128/mBio.00908-16 (2016).

10 Monteiro, J. M. *et al.* Cell shape dynamics during the staphylococcal cell cycle. *Nat Commun* **6**, 8055, doi:10.1038/ncomms9055 (2015).

11 Jorge, A. M., Hoiczyk, E., Gomes, J. P. & Pinho, M. G. EzrA Contributes to the Regulation of Cell Size in *Staphylococcus aureus*. *PLoS ONE* **6**, e27542, doi:10.1371/journal.pone.0027542 (2011).

12 Chen, H., Puhl, H. L., 3rd, Koushik, S. V., Vogel, S. S. & Ikeda, S. R. Measurement of FRET efficiency and ratio of donor to acceptor concentration in living cells. *Biophys J* **91**, L39-41, doi:10.1529/biophysj.106.088773 (2006).

13 Hoppe, A., Christensen, K. & Swanson, J. A. Fluorescence resonance energy transfer-based stoichiometry in living cells. *Biophys J* **83**, 3652-3664, doi:10.1016/S0006-3495(02)75365-4 (2002).
